# A catastrophic marine mortality event caused by a complex algal bloom including the novel brevetoxin producer, *Karenia cristata* (Dinophyceae)

**DOI:** 10.1101/2025.10.31.685766

**Authors:** Shauna A. Murray, Christopher J S Bolch, Steve Brett, Cheong Xin Chan, Mark Doubell, Hazel Farrell, Greta Gaiani, Hannah Greenhough, Gustaaf Hallegraeff, Tamsyne Smith Harding, D. Tim Harwood, Robert G. Hatfield, Neil MacDonald, Ian Moody, Hugo Bastos de Oliveira, Lesley Rhodes, Andrew Selwood, Justin Seymour, Nachshon Siboni, Anastasiia Snigirova, Nikola Streiber, Craig Styan, Anne Rolton, Clinton Wilkinson, Kirsty Smith

## Abstract

Harmful algal blooms of *Karenia brevis* (Dinophyceae) are a global anomaly, occurring in one location worldwide, causing severe marine and acute human impacts via brevetoxins (BTX). During 2025 an unprecedented, currently ongoing, mass marine mortality occurred in South Australia, across an area of ∼20,000 km^2^, persisting for >6 months, resulting in the deaths of ∼10^6^ marine animals of >550 taxa, with human health impacts. Using custom metabarcoding, long-read sequencing and targeted quantitative PCR, we characterized the microalgal assemblage. *Karenia cristata* dominated over the sampling area, in an assemblage with four other *Karenia* species with varied abundances spatially and temporally. High abundances of *K. cristata* appeared in the austral autumn, and hydrodynamic processes appear to have entrained cells coastward in the semi-enclosed seas. We isolated the species and characterized it using light and electron microscopy, liquid chromatography mass spectrometry and toxicity assays. We show for the first time that the rare and little known *K. cristata* produces significant BTX with a profile (BTX-2, -3, -B5), differing from *K. brevis*, with toxicological effects. These findings reveal a novel, significant BTX-producing *Karenia*, which considering its substantial detrimental marine ecosystem impacts, is an emerging international threat with unknown consequences in changing ocean conditions.

## Background

Fish-killing marine harmful algal blooms (HABs), particularly of dinoflagellates (Dinophyceae), are relatively common worldwide, and the vast majority are of a short duration (weeks), with a localized impact^1^. Since March 2025, a marine harmful algal bloom (HAB) off coastal South Australia has spread to encompass an area of ∼20,000 km^2^, resulted in the deaths of millions of individuals of at least 550 marine species^2^. At the time of writing, October 2025, marine mortalities continue^2^. Acute, usually self-limiting human respiratory symptoms have been commonly reported over 6 months in South Australia including the city of Adelaide^3^. Two government inquiries have been established to investigate the causes and consequences of this HAB. Among 849 marine mass mortality incidents recorded in an international database, HAEDAT ^4^ (Supplementary Table 1), this situation ranks among the most impactful of ichthytoxic HABs in marine taxa killed.

In March, a species of *Karenia* (Dinophyceae)*, K. mikimotoi,* was identified as present^5^, and later, brevetoxins were detected in shellfish^3^. Since the mid 20^th^ century^6,7^ it has been recognized that *Karenia brevis* (Dinophyceae) produces potent toxins causing mortalities in marine biota^8^ across trophic levels: zooplankton, invertebrates, fish, birds, marine mammals, in the south-eastern United States^8–10^. In the US, these toxins can become aerosolized and cause respiratory symptoms, as well as accumulate in seafood causing neurotoxic shellfish poisoning in humans ^11,12^. *K. brevis* toxins were characterised as brevetoxins (BTX), lipid soluble polyether compounds that act on voltage-gated sodium channels in vertebrates^13^. *K. brevis* HABs have been a long-standing international anomaly due to: 1) their presence and nearannual recurrence at only one site worldwide, 2) potentially very long durations of HABs (up to 20 months^14^) and 3) severe environmental and acute human health effects^9,12,14–16^. While at least 10 other *Karenia* species, including *K. mikimotoi* can cause large-scale HABs ^15,17–21^, BTX-producing *Karenia* are rare. Those that have been reported, in New Zealand in 1993 ^22,23^, and in the Mediterranean ^24^ have not re-occurred. *Karenia* species other than *K. brevis* producing BTX at a significant concentration^25^ elsewhere have not been identified.

Complicating the investigation into causes of this HAB is that most *Karenia* are cosmopolitan and co-occur ^15^. *Karenia* HABs often consist of 4-5 taxa that differ in toxins produced, ecology and physiology, but are morphologically highly similar or overlapping^17–19,26,27^. Besides BTX, *Karenia* species produce: gymnocin ^28^, gymnodimine ^29^, brevisulcenals, brevisulcatic acids ^30^, brevisin ^31^, brevisamide ^32^, tamulamides ^33^, and other toxins, as well as produce toxic effects due to polyunsaturated fatty acids, superoxide, and unidentified allelochemicals ^1,34–36^. High concentrations of *Karenia* can produce mucilage and anoxia that contribute to marine mortalities^1^. Spatially and temporally distinct *Karenia* assemblages can cause differing impacts, such that the ichthyotoxic effects of a long-running HAB may vary over time and across areas ^37^.

A complex *Karenia* assemblage complicates our understanding of ecological factors contributing to HAB development. Physiology and ecology can differ markedly from one *Karenia* species to another ^15^. *K. brevis*-dominated HABs have been hypothesized to initiate in oligotrophic waters off-shore, followed by coastward entrainment via oceanographic processes ^38^. The extent to which ‘bottom up’ factors, such as increasing temperatures and locally high nutrient coastal inputs are potentially accelerating *Karenia* growth, in comparison to autecological factors and physical concentration effects, is uncertain^38^. In the context of rapidly warming ocean waters^39^ and increasing coastal phytoplankton blooms^40^, the role of changing marine environments in enhancing and intensifying *Karenia* HABs is crucial to establish. Here, we applied a systematic approach to identify and characterize the causative species of the mass marine mortality event, its toxic effects, and factors contributing to this HAB.

## Main

### Environmental and single cell amplification and sequencing

South Australia has a coastline comprising the large semi-enclosed seas of Spencer Gulf (∼320 km long, 116 km maximum width, mean depth 24 m) and Gulf St Vincent∼170 km long, 50 km maximum width, mean depth 21 m), on the south-eastern side of which is the city of Adelaide (Fig. 1a). From March to September 2025, we collected water samples at 39 nearshore and ocean locations (Fig. 1a-c, Supplementary Table 2). Site locations 18-20 in the vicinity of the city of Adelaide (Fig. 1a) were sampled weekly in triplicate from 27 ^th^ July 2025 – 13^th^ September 2025, with weekly sampling continuing at the time of writing. Using a custom, dinoflagellate-specific, long-read MinION (Oxford Nanopore Technologies) sequencing^41^ method focused on regions of the ribosomal RNA gene array on 18 samples, we recovered reads of ∼3020 bp in length from six species belonging to the Kareniaceae family, all with potential fish-killing impacts (Fig. 1d). The Kareniaceae taxa identified were *K. cristata, K. longicanalis, K. mikimotoi, K. papilionacea,* as well as *Karlodinium veneficum* and *Takayama tasmanica* (Fig. 1d). In a phylogenetic analysis of recovered consensus sequences in relation to reference sequences from the NCBI database, each environmental sequence clustered with full support (Fig. 1d) into clades with voucher specimens or multiple identified cultured strains of that taxon (Fig. 1d, Supplementary Table 3).

**Figure 1.**
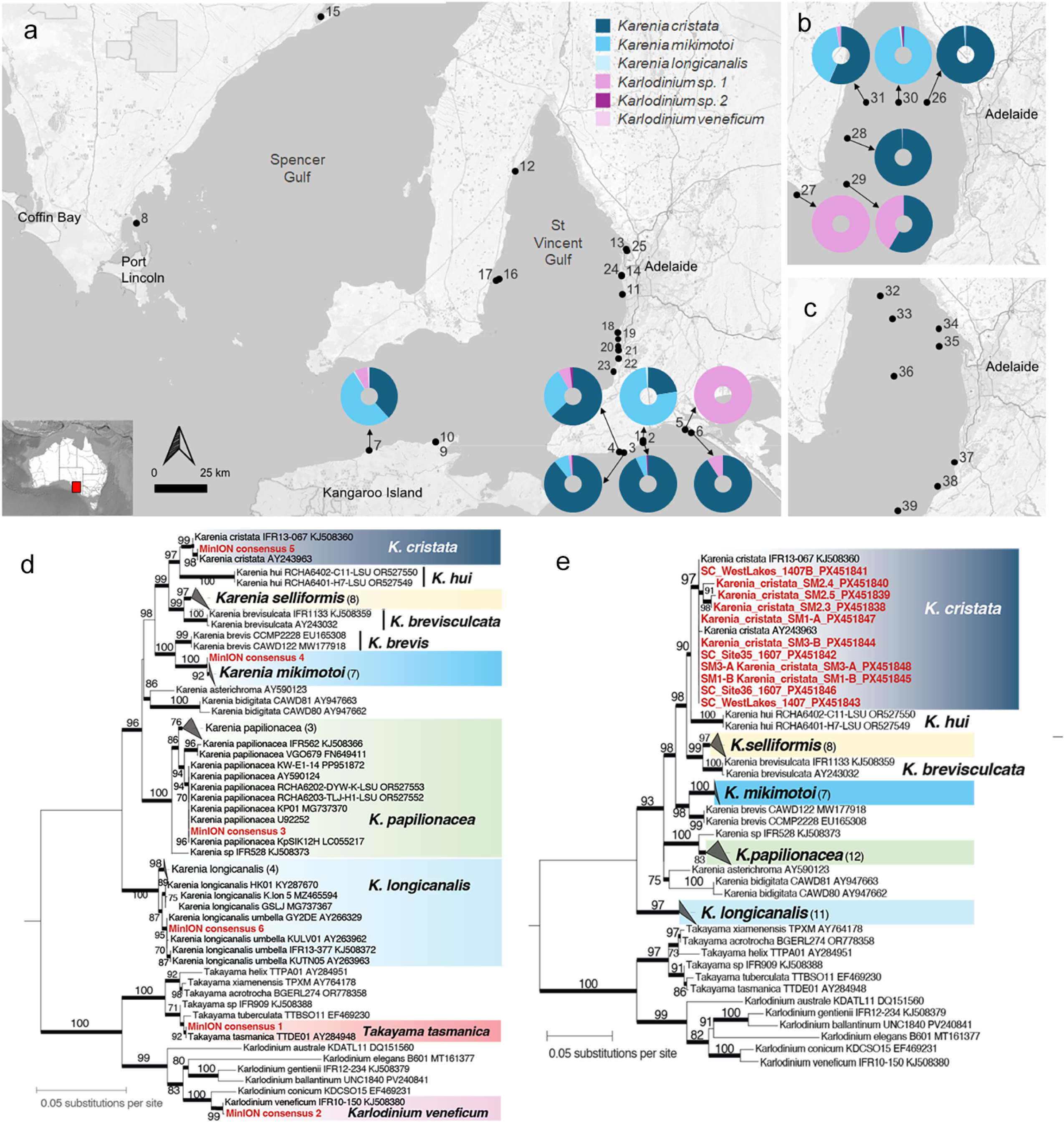
**a-c** Map of South Australian coastline, showing 39 sites at which water was collected (March to September 2025), including two offshore surveys in the St Vincent Gulf. **b.** 22-23^rd^ May and **c.** 16^th^ July 2025. The relevant abundance of ASVs from metabarcoding analyses are shown as donut plots, highlighting the mixed presence of several *Karenia* species and *Karlodinium*. **d, e**. Maximum likelihood tree of Kareniaceae based on LSU rDNA sequences, incorporating **d**. MinION amplicon sequences and **e.** single-cell qPCR amplified sequences generated from this study. Each tree is rooted with the clade of *Takayama* and *Karlodinium* as outgroup to *Karenia.* Sequences generated from this study are highlighted in red. Ultrafast bootstrap support (>70%) is shown on each internal node, and a thick branch length indicates Bayesian posterior probability = 1.0 in the additionally inferred Bayesian tree (see Methods). Unit of branch length is number of substitutions per site.

We used a custom-dinoflagellate Illumina metabarcoding sequencing^42^ method to select amplicon sequence variants (ASVs) identified as Kareniaceae species from 13 marine samples (Fig. 1a, b, Supplementary Tables 2 and 4). We recovered ASVs with high affinity to the species *K. cristata, K. longicanalis*, *K. mikimotoi, Karl. veneficum* and two unidentified *Karlodinium* spp (Fig. 1a, b, Supplementary Table 4). The proportions of the taxa present varied by sample, with the most abundant kareniacean taxa being *K. cristata, K. mikimotoi,* and an unidentified *Karlodinium sp* (Fig. 1a, b). *K. cristata* dominated at sites 2-4, 6, 26, 28 and 31. *K.mikimotoi* was abundant and dominant in three samples (1, 7, and 30, Fig. 1a, b), in line with the identification of that taxon based on light microscopical observation in samples from Victor Harbor (sites 1-4) ^5^. At sites 5, 27, no *Karenia* were found (Fig. 1a, b). Single cell picking, followed by direct PCR and sequencing of an rRNA barcoding region (partial 28S rRNA) was conducted on cells from samples from sites 24, 35 and 36, and the sequences were found to be *Karenia cristata* (Fig 1e, samples labelled SC).

### Species-specific *Karenia* quantification across 39 locations from March to September 2025

Following the identification of *Karenia* using high-throughput sequencing, we proceeded to characterize assemblages via quantitative PCR (qPCR), using published assays^43^ for *K. mikimotoi, K. longicanalis,* and *K. papilionacea* (Fig. 2, Supplementary Table 5). As critical *Karenia* diversity may not have been present in initial sequenced samples, we additionally tested for *K. brevisulcata*^43^, and *K. brevis*^44^. We designed assays for *K. cristata* and *K. hui* (Fig. 2, Supplementary Fig. 1), tested them against a set of multiple strains of seven *Karenia* species (Supplementary Tables 3, 5) and found no cross-reactivity. Assays were highly efficient (Supplementary Fig 1). Across sites and months, we detected five species of *Karenia: K. cristata, K. mikimotoi, K. brevisulcata, K. longicanalis* and *K. papilionacea*, at concentrations of 1-10^6^ cells L^-1^ (Fig. 2a). No *Karenia brevis* or *K. hui* were detected. This showed us that *K, brevis*, the only known high BTX producing species, was not present and not a toxin source. *Karenia hui* is morphologically and genetically highly similar to *K. cristata*^18^ (Fig 1d, e), and establishing its absence in this HAB was important. Of *Karenia* species present, approximately 90% of samples contained a majority of *K. cristata* compared to other *Karenia* spp, at abundances of 10^3^ - 10^6^ cells L^-1^, with lower levels of *K. mikimotoi* ∼10^3^ cells L^-1^ present (Fig. 2). *K. brevisulcata,* which had not been identified in the initial samples using metabarcoding or long read sequencing, was found (Fig. 2). Regional differences in *Karenia* assemblages and abundances were apparent. Samples from Spencer Gulf in June and July (sites 8, 15, Fig. 1a) and the western St Vincent Gulf (sites 12, 16, 17) were occasionally dominated by *K. papilionacea* (Fig. 2). These samples also showed that *Karenia* could aggregate or accumulate in exceptionally dense layers in the water column. A surface slick showing a maximum concentration of 15 x 10^6^ cells L^-1^ of *K. cristata* was found in one sample from July (site 17). Weekly samples from the sites around metropolitan Adelaide showed a consistent, high abundance of *K. cristata* from July-September, of ∼10^3^ - 10^6^ cells L^-1^, with lower levels of *K. mikimotoi* ∼10^3^ cells L^-1^ present, and trace *K. brevisulcata*, with little sign of increasing or decreasing trends over the period (Fig. 2).

**Figure 2.**
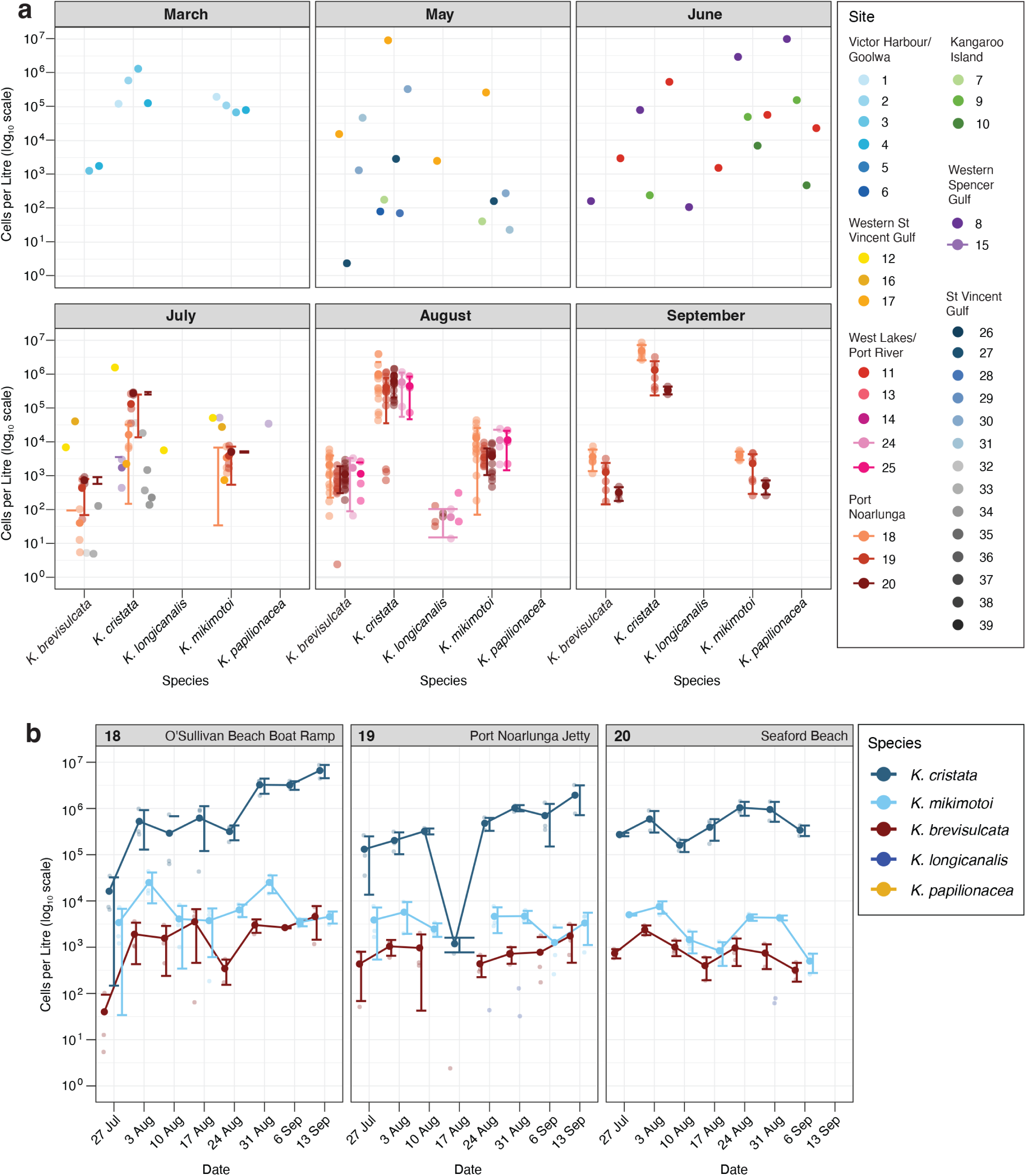
Cell abundance of distinct *Karenia* species detected in South Australian waters throughout the period of March through September 2025, based on qPCR detection. **a**. Cells L^-1^ (in log10 scale) is shown for samples collected at 35 distinct sites in March, May, June, July, August, and September 2025, for each of the detected species of *K. brevisculcata*, *K. cristata*, *K. longicanalis*, *K. mikimotoi*, and *K. papilionacea*. *K. brevis* and *K. hui* were not detected in any samples. The sites were colour coded and grouped by the seven regions as shown in the legend key. Error bar indicating standard deviation is shown for site with two or more samples. **b**. Cells L^-1^ (in log10 scale) is shown for weekly samples collected at the O’Sullivan Beach Boat Ramp (site 18), Port Noarlunga Jetty (site 19) and Seaford Beach (site 20) in the vicinity of Port Noarlunga, between 27 July and 13 September 2025. Data are shown for each detected *Karenia* species across the 8 sampling occasions during the period. Mean and error bar indicating standard deviation are shown for triplicate samples where available.

### *Karenia cristata* isolation into culture and morphological characterization

In early May 2025, shellfish farms at the western Gulf St Vincent and Kangaroo Island (Fig 1a) were closed due to the detection of BTX^3^. These were the first ever BTX detections in Australia. Hence, to characterize *Karenia* BTX producers, we isolated single cells into a seawater media in laboratory culture. In particular, we focused on *K. cristata* due to its ubiquity across locations (Fig. 1a-e). From sample 6 collected on 7^th^ May (Fig. 1a), three non-clonal, non-axenic cultures (SM1, SM2, and SM3) were isolated. DNA was extracted and PCR was used to amplify a dinoflagellate barcoding gene (partial 28S rRNA). In the phylogenetic analysis of the resulting sequences (Fig. 1e), the cultured strains (SM1, SM2, and SM3) grouped with high support to a clade comprising the only two verified sequences of *Karenia cristata* isolated, respectively, from South Africa^45^ and a French island south of Newfoundland, Canada ^46^.

Morphological investigations were undertaken on strain SM2 as well as *Karenia* cells from samples dominated by *K. cristata* (Fig. 3) using light and electron microscopy. *K. cristata* cells were fragile, fast swimming, dorso-ventrally compressed (Fig. 3), with numerous yellow-green irregularly rod-shaped chloroplasts to spherical (Fig, 3a-c, f), a short epicone with hemispherical outline, a longer asymmetrical bi-lobed hypocone with the right lobe slightly longer than the left (Fig. 3c, g, h). Cingular displacement was ∼28% of cell length (Fig. 3b).□Cell length ranged from 21.0 to 32.2µm (mean = 24.8 µm, SD=2.32, n=40), 16.1 to 28.6 µm wide (mean=21.1 µm, SD=2.67; n=40), and 9.1-13.6 µm depth (mean=11.2 µm, SD=1.35, n=20). The length to width ratio of cells ranged from 1.06 to 1.22 (mean=1.14, SD=0.04). Living cells exhibited a straight apical groove that was elevated on the ventral side to form a small apical crest. (cr; Figs. 3b, c). The crest became less evident when cells were stressed or immobilised under a coverslip. Cells examined using scanning electron microscopy (SEM) showed the crest was formed by the right side of the apical groove being slightly raised (Figs. 3g-k). After Lugol’s iodine preservation, some cells exhibited a more pointed epicone and a more pronounced apical crest, with the nucleus positioned lower in the hypocone (Fig. 3d). A straight apical groove extended on the ventral surface of the epicone, with a narrow depression (fe; Fig. 3h) extending dorsally to the hook-shaped ventral flange (vf; Figs. 3a, g). On the dorsal side, the apical groove extended to ∼1/3 of epicone length (ag; Fig. 3i). Using DIC light microscopy, the sulcal intrusion (si) into the epicone appeared closed or slightly open (si, Figs. 3a, b), but SEM elucidated the sulcal intrusion to be either wide-open or end in a finger-like projection (Figs. 3g, h). Comparisons of morphological features to other similar *Karenia* species is given in Supplementary information 1.

**Figure 3.**
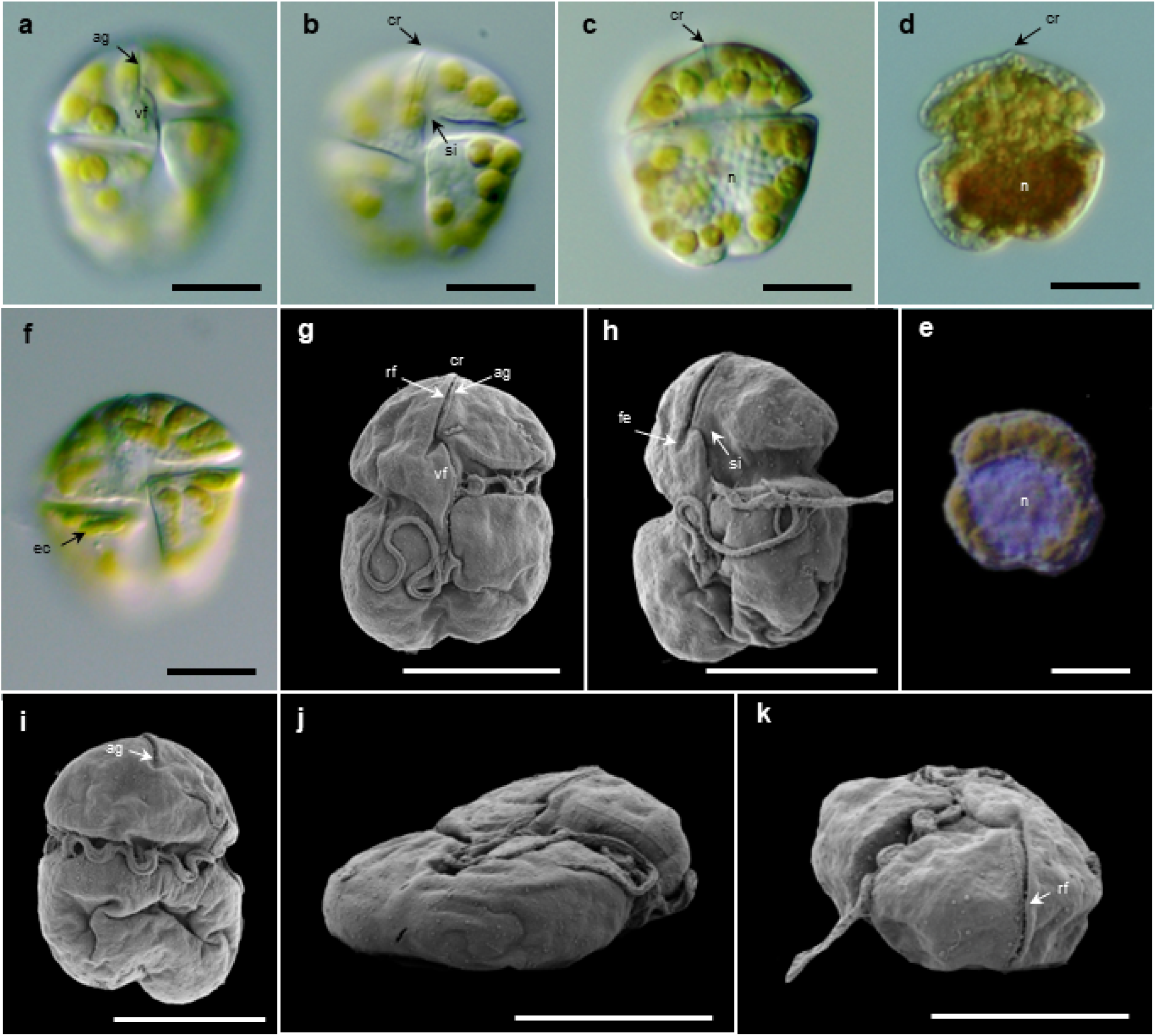
Morphology of *Karenia cristata* from South Australian blooms and laboratory cultures. **a-f.** LM (DIC) of *K. cristata* from blooms. **a-b**. Live cells in ventral view showing key features including the apical groove (ag), apical crest (cr), sulcal intrusion (si) and ventral flange (vf). **c**. Live cell dorsal view (n=nucleus). **d**. Lugol’s fixed bloom cell (n=nucleus) with a more prominent apical crest (cr). **e**. Composite DIC/FM image of a Lugol’s fixed cell from *K. cristata* culture SM2; lugol’s fixed cell image overlaid by FM image of cell after DAPI staining showing the circular to ovoid outline of the nucleus (n). **f.** Ventral view live cell with elongated irregular rod-like chloroplasts (ec). **g-k**. SEM of *K. cristata* SM2 cultured cells. **g**. Ventral view with key surface features including the apical crest (cr) formed by a raised flange (rf) on right edge of apical groove (ag). **h**. Ventral-lateral view showing sulcal intrusion (si) into the epicone and a finger-like depression (fe) extending from apical groove. **i**. Dorsal view showing apical groove extending approx. 1/3 down dorsal side of the epicone. **j**. Antapico-lateral view showing dorso-ventral compression of cells. **k**. Apico-lateral view showing straight apical groove with raised flange (rf), extending over apex of cell.

### Toxicology and toxicity

To examine the production of BTX by strains of *K. cristata*, an aliquot of dense culture of SM2 was extracted and examined using liquid chromatography tandem mass spectrometry (LC-MS/MS) against verified BTX reference standards for 8 analogs (Supplementary Table 7). Results showed a high concentration of the BTX analogues BTX-2, BTX-3 and smaller amounts of BTX-B5 (Fig. 4a, b) in strain SM2 and in two samples, with an absence of other analogs at detection levels.

**Figure 4.**
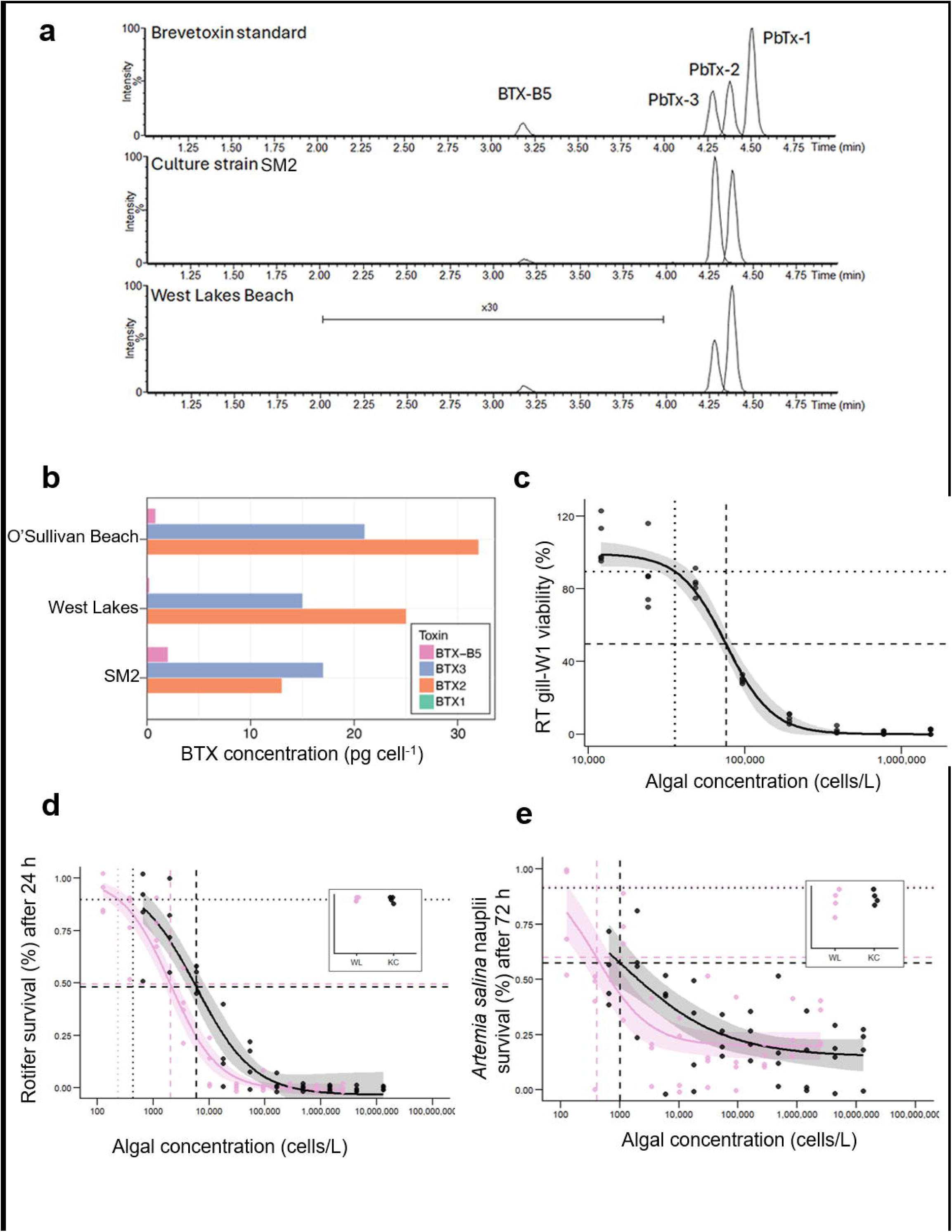
Toxins present and toxicological impact on a fish gill cell line. **a.** LC-MS chromatogram showing BTX standards, and BTXs detected in a seawater sample collected during the 2025 South Australia bloom, and from a *Karenia* culture derived from the same bloom event; **b**. Quantification of brevetoxin analogues per cell, illustrating similarity in toxin profiles and cellular production rates. **c**. Rainbow trout fish gill cell viability assay showing the percentage mortality in relation to differing concentrations of a water sample from West Lakes on 6^th^ September 2025. Horizontal dashed and dotted lines indicate 50% and 10% survival, with corresponding vertical dashed and dotted lines showing LC50 and LC10 estimates, respectively. **d** Proportion of *Brachionus plicatilis* rotifers surviving after 24 h exposure to increasing concentrations of *K. cristata* cells from a laboratory culture (SM2; black) and a field-collected West Lakes bloom (pink). **e** Proportion of *Artemia salina* nauplii surviving after 72 h exposure to the same algal sources. Points show observed proportions per replicate well; solid lines indicate fitted dose–response models with shaded 95% confidence intervals. Horizontal dashed lines indicate 50% survival, with corresponding vertical dashed lines showing LC50 estimates for each algal source. Insets show survival proportions in control wells (artificial seawater only).

To further examine *K. cristata* ichthyotoxicity, a Rainbow Trout (*Oncorhynchus mykiss*) gill cell viability assay was conducted using supernatant of the same *K. cristata* strain (SM2)^47,48^. Exposure of RTgill-W1 cells for only 2 h to *K. cristata* SM2 supernatant resulted in an LC50 estimated at 75,625 ± 4597 cells L^-1^ and an LC10 of 35851 ± 6017 (Fig. 4c).

The strain SM2 and parallel assays of a sample of the bloom collected from West Lakes on 14th January 2026 were tested with whole-organism mortality assays using rotifers (*Brachionus plicatilis*) and brine shrimp (*Artemia salina*) nauplii over 24 and 72 h, respectively. Rotifers had a low LC50 after 24 h exposure (SM2: 5845 ± 1688 cells L□¹; West Lakes bloom: 2032 ± 313 cells L□¹), with a small but statistically significant difference between algal sources (t = 2.22, df = 14, p = 0.04); corresponding LC10 values were 432 ± 170 cells L□¹ for SM2 and 235 ± 61 cells L□¹ for the West Lakes bloom (Fig. 4d). *Artemia* nauplii showed slower and incomplete mortality over 72 h, but displayed clear concentration–response relationships, with no detectable difference in LC50 values between algal sources (SM2: 1012 ± 556 cells L□¹; West Lakes: 406 ± 132 cells L□¹; t = 1.06, df = 14, p = 0.30), and LC10 values of 12 ± 25 cells L□¹ and 41 ± 27 cells L□¹, respectively (Fig. 4e), noting that LC10 estimates required extrapolation beyond the lowest tested concentrations (800 cells L□¹ for SM2; 150 cells L□¹ for the West Lakes bloom). Mean measured algal concentrations remaining in the highest concentration wells were at 87% of nominal levels at the end of the Artemia exposures.

### Chlorophyll a anomaly and temperatures over the period

We used the remotely sensed chlorophyll-a concentration anomaly, as compared to a 21 year climatology (2002-2023) (Fig. 5a) to determine departures from this long-term daily climatology based on the previous 10 day median filter to show the potential distribution of *Karenia* species throughout the region. This method shows that chlorophyll a rose sharply during the study period in relation to previous years, reaching concentrations well above the typical autumn baseline (Fig. 5a). Phytoplankton biomass expanded northwestward in autumn through the Investigator Strait and into western Gulf St Vincent (Figure 5b), reaching the Adelaide metropolitan coast in winter (Figure5c). Elevated biomass has persisted for several months, with highly abundant *Karenia* species observed even in deeper areas of the Gulf (∼30m, Fig. 5 b,c). Throughout the period, temperatures in the area of the HAB in the austral autumn varied from around 14 – 21 °C (Fig 5d). During May – September, when *Karenia* cells were most abundant and widespread, temperatures were around 14 – 18 °C (Fig 5d).

**Figure 5.**
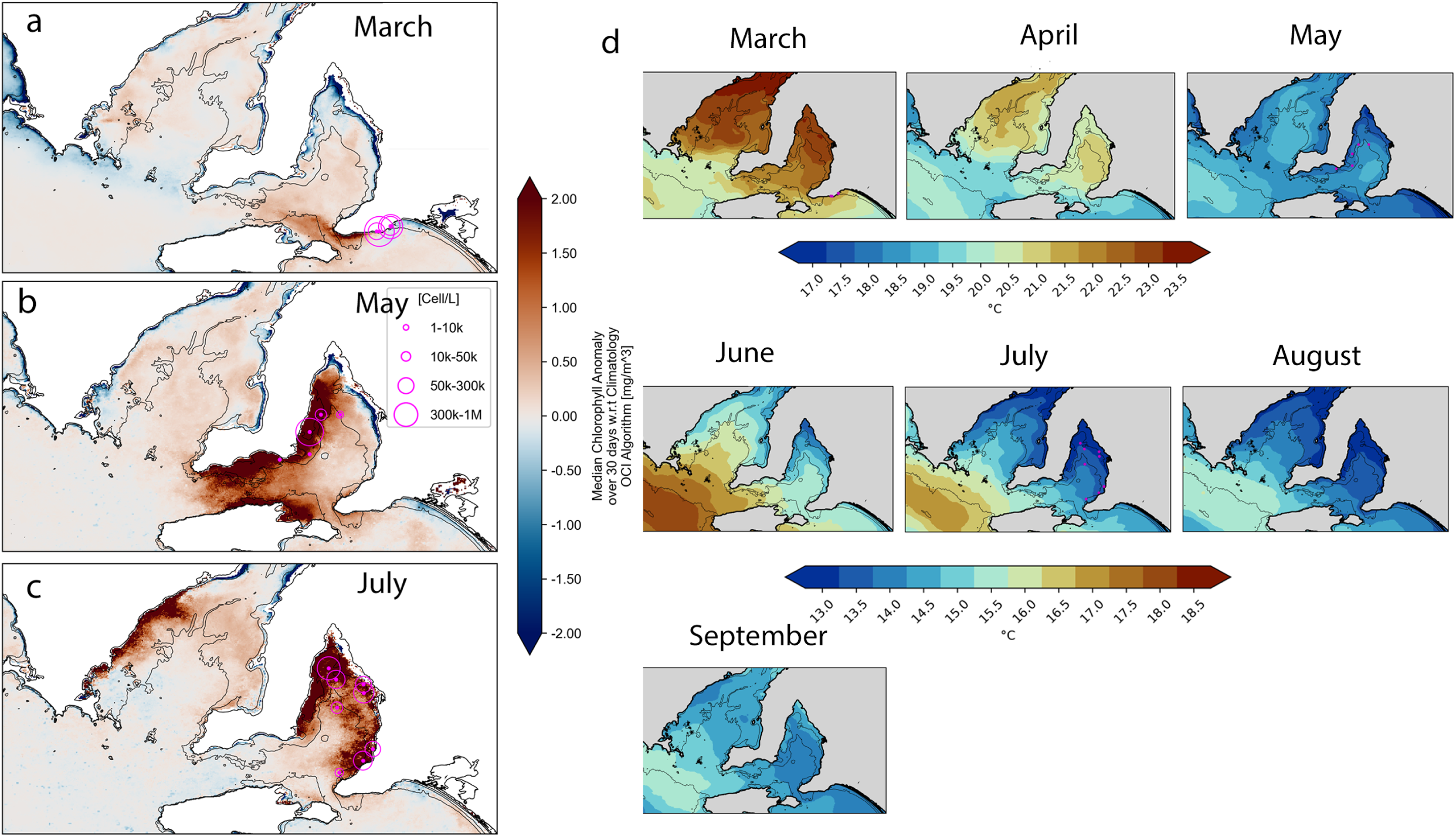
Monthly maps (pseudo-colour) of Chlorophyll-A anomalies off the South Australian coastline based on a 21 year dataset, and temperature across the period. **a**. March mean anomalies (stations 1-4), **b.** May (with stations 27-31), **c**. July (stations 32-39). The anomalies are computed with relation to a 21 year climatology (2002-2023) from MODIS and VIIRS sensors (see Methods) using the OCI algorithm. The magenta circles represent the total abundance of *Karenia* spp in the selected samples, while the light black line represents the 10m and 30m isobaths. **d.**Temperatures in the area from March to September 2025, during the HAB,

## Discussion

Here, we describe the catastrophic effects of a high brevetoxin producing marine dinoflagellate, *Karenia cristata*. Our analysis reveals that the *K*. *cristata* is the second identified copious Kareniaceae brevetoxin producer after *K. brevis*, a species which has a restricted distribution in the south-eastern United States^15^. *K.cristata* co-occurred in a multi-species kareniacean assemblage with four other species including *K. mikimotoi,* leading to a HAB that has, at the time of writing, lasted over >6 months, spread across an area of ∼20,000 km^2^ (Figs 1, 2), and resulted in devastating impacts on marine animals, as well as acute human health impacts. While marine ichthyotoxic HAB events can be common worldwide, including in Australia, of 849 reported events from 1985-2025^4^ (http://haedat.iode.org), this is amongst the most destructive and widespread marine ichthyotoxic HAB, in terms of temporal and spatial extent and particularly in the diversity of marine taxa killed. Such wide ecosystem and human health effects of ichthyotoxic HABs are most pronounced for species producing known toxins ^1^, such as some of the Karenianceae (Supplementary Table 1), rather than raphidophytes (ie *Heterosigma*) and dictyochophytes (ie *Pseudochattonella*) that require surface contact between the microalga and marine animals for harmful effects ^1^.

We show that a strain of *Karenia cristata* that we isolated and cultured from the impacted region produced three analogues of BTXs in similar (Fig. 4, ∼30-40 pg cell^-1^) quantities to *K. brevis* cultures (∼10-20 pg cell^-1^ ^23^ ^49^). *K. brevis* produces brevetoxins with type A and type B polyether backbones, with BTX-1 (BTX-1) and BTX-2 (BTX-2) ^13,50^ the respective parent structures, with other known brevetoxins as structural derivatives of these core compounds^51^. While strains of *K. brevis* typically produce approximately 30% BTX-1, here, we found a notable absence of type A polyether backbones including BTX-1, marking a distinctive toxin profile for *K. cristata* (Fig. 4). Reports of brevetoxins not associated with *K. brevis* are very rare. A strain of *K. papilionacea* was reported to produce brevetoxins, albeit at fg levels ^25^. Corsica in the Mediterranean Sea reported BTXs in mussels once ^24^. While no *K. cristata* was found at the time^24^, given the rarity of BTX production in association with *Karenia* outside of the south-eastern United States, it may be possible that *K. cristata* was present and unrecognised. In New Zealand in 1993, a bloom of *Karenia* spp. occurred in the Hauraki Gulf and only type B BTXs were detected in shellfish^23^, indicating that the producing microalgal species, which was not identified, likely produced only type B BTXs. *K. cristata* may have been responsible for this Neurotoxic Shellfish Poisoning event.

*K. cristata* has only ever been previously reported from two locations internationally: False Bay and Walker Bay, South Africa, in the mid-1990s, and a remote island off the coast of Newfoundland (French Territory) ^46^. In South Africa, *K. cristata* caused an ichthyotoxic HAB (Supplementary Table 1), causing large mortalities of marine life in March 1989 ^52^ and in the same region in 1995-1996 ^45,53^. Human health symptoms included eye, nose, throat and skin irritations, the exact cause of which was not further investigated. In HABs caused by *K. brevis*, brevetoxins commonly cause skin irritations, and can become aerosolised as the fragile cells break and cause respiratory distress in humans, particularly during onshore winds ^54^ ^11,12^. The US Center for Disease Control (CDC) states that inhaling brevetoxins can lead to serious health effects including: shortness of breath, asthma exacerbation, bronchoconstriction, bronchitis pneumonia (https://www.cdc.gov/harmful-algal-blooms/hcp/clinical-signs/symptoms-saltwater-harmful-algal-blooms.html), some of which have been very commonly reported in South Australia since March. Hence it seems likely that the reported human respiratory symptoms both in this event and the South African *K. cristata* HAB are caused by brevetoxins.

Understanding the sources of the ichthyotoxicity and marine animal mortalities in this event is complex, due to multiple Kareniacaea co-occurring and varying in abundance regionally (Figs 1-2, 4). Nonetheless, determining toxicological source is critical to decision-making regarding actions for mitigation, prevention or restoration of marine taxa, as methods may be more or less suitable depending on what exactly caused the marine mortalities. Some toxins such as brevetoxins persist environmentally for some time in the absence of *Karenia* cells, while other toxic effects are more volatile or disappear rapidly at cell death. Brevetoxins or other characterised toxins, on the other hand, may not be of high importance in causing ichythotoxicity, as pure brevetoxins may be only weakly toxic in fish gill bioassays ^35, 55^. Besides *K. cristata,* the species *K. mikimotoi, K. brevisulcata, K. longicanalis* and *K. papilionacea* were present and occasionally dominated (Fig 2), and have documented ichthyotoxic effects, as reported internationally^17,18,27^. Similarly *Karlodinium veneficum* and *Takayama tasmanica* (Fig. 1, 2) have caused ichthyotoxic HABs ^48^. In this sense, our results can are the starting point rather than the end in investigating the degree to which well-characterised toxins such as brevetoxins, as opposed to 1) toxins: gymnocin ^28^, gymnodimine ^29^, brevisulcenals, brevisulcatic acids ^30^, brevisin ^31^, brevisamide ^32^, tamulamides ^33^, and 2) compounds other than chemically characterised toxins, including polyunsaturated fatty acids, superoxide, and multiple unidentified allelochemicals ^1,34–36^ all contributed regionally or over time to the observed marine mortalities. *Karenia* species were also extremely abundant in the water column (10^6^ cells L^-1^) at some locations, in which case marine mortalities may have occurred simply due to anoxia ^1^.

Investigations of Australian *Karenia* diversity are important to determine whether *K. cristata* has long been present, or whether populations have established relatively recently, through processes such as ballast water exchange. *Karenia* blooms have occurred in Australia^26,56,57^ but it has also been known for the past two decades that undescribed *Karenia* species exist in Australian waters ^26,56^. Two previous HABs identified as *Karenia mikimotoi* affected Coffin Bay in March 1995 and February-March 2014, and resulted in deaths of fish and invertebrates ^58^. A HAB of *K. longicanalis*^56^ caused extensive fish kills in Tasmania. Australia has previously recorded 6 species of *Karenia,*7 species of *Karlodinium* and 3 species of *Takayama,* most of which are capable of ichthyotoxicity (Supplementary Fig. 2). In re-investigating the source of fish deaths in the 2014 South Australian (Coffin Bay) HAB using our environmental DNA samples^57^ (Supplementary Fig 2), we confirmed that *K. mikimotoi* was the only *Karenia* species present at the time. Our inability to determine whether *K. cristata* has long been present or not in Australian waters highlights an urgent need to investigate marine microbial diversity. Such studies need to use methods that allow for an understanding of the ‘hidden flora’ or rare species at low abundances, to establish baselines and to prepare for the changes we are experiencing in marine ecosystems.

The environmental drivers behind this unprecedented HAB are unresolved, but may be multifarious and linked to anomalous oceanographic conditions. The onset of the bloom closely coincided with a temporally extensive and periodically severe marine heatwave (MHW) in this region^59^, with water temperature anomalies in the range of 2-3° C (Fig. 5). Globally, increased frequency of HABs has been linked to rising seawater temperatures^60^. An unprecedented HAB caused by another *Karenia* species, *K. selliformis*, was preceded by a significant MHW in Japan^61^. However, different *Karenia* species exhibit distinct temperature and nutrient preferences^15^ and the temperature for optimum growth of *K. cristata* has not been defined. The only other previously reported HAB of *K. cristata* took place in South Africa during the austral autumn in relatively cool water temperatures of 14-17 °C ^45,52^. This pattern is consistent with the expansion of the HAB in South Australian waters after May 2025 (Fig. 5d), as the highest *Karenia* abundances coincided with lower temperature conditions. Therefore, it is possible that the timing of bloom onset was unrelated to the concurrent MHW, or was triggered by other physico-chemical conditions linked to the MHW (e.g. heightened water column stability, increased bacterial nutrient cycling). The relatively rapid and widespread geographic expansion of this event bears some of the hallmarks of interannual blooms of other *Karenia* species, which often build slowly offshore in oligotrophic waters and spread into coastal waters in high concentrations via hydrodynamic processes ^38^. Temporal changes in remotely sensed chlorophyll-a anomalies during the HAB expansion across St Vincent and Spencer Gulfs (Fig, 5) align with the net seasonal circulation patterns in the region, which is governed by both wind and density-driven processes, and can entrain cells coastward in these highly saline (inverse) estuaries characterized by high evaporation and low river inputs and precipitation ^62, 62, 63, 64^.

Beyond the physical concentration effects of oceanographic processes, there remains the question of why this bloom has been characterised by such significant spatial expansion and temporal persistence. There is the potential that autecological processes, acting in synergy with oceanographic triggers, could be at play. For instance, several *Karenia* species show allelopathic capacity^65, 34,66,67^, allowing them to kill or inhibit co-occurring phytoplankton or predators, leading to their domination of the micro-algal community. Laboratory experiments of *K. cristata* physiological ecology would be necessary to establish these factors. The sudden emergence of *K. cristata* as a potentially new cause of significant coastal HABs, specifically, only the second known *Karenia* HAB producing unprecedented concentrations of highly damaging brevetoxins, has occurred in the context of evidence that climate change has expanded conditions favourable for HABs worldwide ^60, 40,68^,. Given that *K. cristata* has a distribution in the North and South Atlantic Ocean as well as the Great Australian Bight, and could potentially impact many countries with suitable conditions, this highlights an urgent new need to develop a solid mechanistic understanding of the triggers of the development and persistence of *Karenia cristata* HABs within multispecies Kareniaceae assemblages with varying toxicologies. Such an effort is critical to establish early warning and mitigation strategies to protect environments and human health from brevetoxin exposure.

## Methods

### Water sample collection

Water samples were collected between 18 March and 13 September 2025 from 39 sites off the South Australian coast (Supplementary Table 2, Fig. 1a-c), from nearshore locations (18^th^ March-13^th^ September 2025, Fig. 1a) and on two occasions samples at sea, onboard SARDI’s RV Ngerin (22^nd^-23^rd^ May 2025, Fig. 1b) and FPV Southern Ranger (16 July 2025, Fig. 1c). Locations were presented (Fig. 1a-c) using QGIS ^69^, and an Australian national basemap from Geoscience Australia. Where indicated (Supplementary Table 2), surface samples were a direct sample within the top 0.5 m of the water column either collected by boat or from the jetty or shore. Three samples were collected via a 25cm integrated tube sampler (4m). Sub-surface samples were collected via a 5 L Niskin bottle from 1m or at the deep chlorophyll maximum (DCM), as determined by CTD cast using a SBE 19+ CTD profiler (Sea-Bird Scientific).

Unpreserved (live) samples were stored cool, in the dark and transported as soon as possible to the laboratory at ambient temperature. Sub samples (0.1-1 L) of live samples were filtered on 5 µm Durapore (Merck, Millipore) filters using a vacuum pump and held at −20 °C until further processing. Samples indicated by ^ in Supplementary Table 2 were fixed with acidified Lugol’s solution (∼5ml/L) and stored in the dark before filtration. At locations indicated by * in Supplementary Table 2, water samples were collected on filters and preserved in Longmire’s buffer using an onsite gravity filtration method^81^.

### Remote sensing

Deeper waters (>10 m) were examined using chlorophyll-A IMOS Surface Remote Sensing products using the OCI Algorithm^82^ and quality control. We merged both Visible Infrared Imaging Radiometer Suite (VIIRS) abord the NOAA20 and SNPP satellites, and use the Moderate Resolution Imaging Spectroradiometer (MODIS) to compute a 21 year climatology (2002-2023), via LombScargle Periodogram^83,84^. Anomalies (Fig. 5a-c) are departures from this long-term daily climatology. Intermittent blooms were characterised by a previous 10 day median filter, due to cloud cover. Although estimates in very shallow water (<15 m) are in generally highly biased and noisy, the signal increases detected in shallow waters were much higher than any previous estimates (not shown).

### Microscope observation

Light microscopic examinations were undertaken on live and Lugol’s preserved cells using Zeiss (Axiolab A1, Axioscope A1 and Axioplan 2 Imaging) upright compound microscopes equipped for bright field, phase-contrast, DIC and epifluorescence microscopy. Samples were examined directly as wet mounts on slides, or after gentle concentration using gravity-assisted membrane filtration. Cell counts were conducted in Sedgewick-Rafter Counting cells. The abundance of cells in culture was established by counting subsamples preserved in Lugol’s Solution using Utermöhl chambers and an inverted microscope (Olympus CK41 microscope). Cell abundance was used to calculate the number of cells in the DNA extractions to generate cell based standard curves for quantitative PCR analyses.

For scanning electron microscopy, live cells in seawater were fixed (2h) with freshly opened 1-2% osmium tetroxide, washed 3 times with distilled water, followed by a dehydration series of ethanol (30, 50, 70, 80, 95, 100%, 10 min each), then chemically dried first with a 1:1 mixture of hexamethyldisilazane (HMDS):ethanol, followed by 100% HMDS (10 min each). Similar fixation protocols starting with Lugol or 1-4% glutaraldehyde preserved cells consistently failed to produce satisfactory results. Cells were collected on nucleopore filters, mounted on aluminium stubs, coated with platinum (5nm layer thickness), and examined with a Hitachi SU70 field emission scanning electron microscope (FESEM).

### Culture methods

Live cells of *Karenia* were selected from samples enriched by addition of K medium ^70^ using flame-drawn Pasteur pipettes, using either a Nikon Inverted light microscope or a Leica MZ9.5 brightfield/darkfield stereomicroscope. To establish cultures, groups of 10-50 *Karenia* cells were drawn up from K/2 enrichment samples using a sterile 200 ul pipette/tip, transferred to a 12-well plate, and incubated at 17 C and 12:12 light (80 umoles PAR m-2s-2) for 1-3 days. After incubation, over successive weeks, any non-target species were carefully removed by micro-pipette, and plates returned to the same incubation conditions. From samples collected in May 2025 at site 6 (Fig. 1a), three non-clonal, non-axenic cultures (SM1, SM2, and SM3) were isolated and maintained.

### DNA extraction

DNA was extracted from water samples and cultures using the DNeasy 96 PowerSoil Pro QIAcube HT Kit (Qiagen) following the manufacturer’s instructions and stored at –20 □until further analysis. Nineteen *Karenia* and *Karlodinium* cultures from the Cawthron Institute Culture Collection, New Zealand, were used as standards (Supplementary Table 3). Set volumes of culture were filtered onto 0.45 μm Durapore® PVDF Membrane filter (Sigma-Aldrich) and DNA was extracted from filters as described above.

### qPCR analysis

All qPCR assays were prepared using the epMotion 5075I Automated Liquid Handling System and run on a Bio-Rad CFX384 Touch Real-Time PCR Detection System with CFX Maestro software v2.3. Each assay included three technical replicates, a standard curve, and negative controls. Standard curves were generated using serial dilutions of DNA extracted from known cell numbers of *Karenia* and *Karlodinium* species maintained in culture (Supplementary Table 3). The reaction mixture for each assay consisted of 2.5 µL of iTaq Universal Probes Supermix (Bio-Rad), variable concentrations of forward and reverse primers and probe (Supplementary Table 5), 1 µL of template DNA, and sterile water to a final volume of 5 µL. Thermal cycling conditions were as follows: initial denaturation at 95□°C for 3 minutes, followed by up to 45 cycles of 95□°C for 15 seconds and 60□°C for 1 minute. For quality control, the coefficient of variation (%CV) was calculated for the Cq values of technical triplicates. If the %CV exceeded 2%, one replicate was excluded. Biological samples with a %CV greater than 5% were omitted from further analysis. Final data were normalized to cell copies per liter of seawater.

### Single-cell qPCR

For the identification of individual cells of *Karenia* species present in samples, swimming live cells were picked individually from samples using a fine glass Pasteur Pipette, using a Nikon Inverted light microscope. Cells were individually photographed under 40 x magnification, then placed in PCR tubes, and frozen at −80 degrees. PCR reagents were added directly to tubes.

### Illumina metabarcoding

The 28S ribosomal RNA (rRNA) gene was amplified from select environmental samples using primers targeting the D1-D2 region of the 28S rRNA and included the tails for NEXTERA sequencing: D1R-F (5’-ACC CGC TGA ATT TAA GCA TA-3’) ^71^ and 305-R (5’-TTT AAY TCT CTT TYC AAA GTC C-3’) ^42^. PCR reactions were undertaken with 450 nM of each primer, 12.5 µL of 2 × MyFi™ Mix (Bioline), approximately 5 ng of DNA and sterile water for a total reaction volume of 25 µL ^42^. Cycling conditions were: 95 °C for 5 min, followed by 35 cycles of 94 °C for 30 s, 54 °C for 30 s, 72 °C for 45 s, and a final extension of 72 °C for 7 min. PCR products were pooled and visualised on 1.5 % agarose gel with Red Safe™ DNA Loading Dye (Herogen Biotech) and ultraviolet illumination. PCR negatives were run to assess for contamination during the PCR steps. The PCR products were purified, cleaned of primer dimers, and normalised using SequalPrep™ Normalization Plate (ThermoFisher), and submitted to Sequench Ltd. (Nelson, Aotearoa-New Zealand) for library preparation.

For the bioinformatic analysis, raw reads were processed, after primers were removed with cutadapt ^72^, using the DADA2 package ^73^ within R. Reads were truncated to 200 base pairs (bp) and filtered with a maxEE (maximum number of ‘expected errors’) of 2 for both forward and reverse reads (reads not reaching this threshold were discarded). DADA2 constructs a parametric error matrix (based on the first 108 bp in the dataset), dereplicates the samples, and sequence variants for the forward and reverse reads are inferred based on the derived error profiles from the samples. Singletons observed in the inference step are discarded. Subsequently, paired-end reads were merged with a maximum mismatch of 1 bp and a required minimum overlap of 10 bp. Forward and reverse reads that did not merge were not included in further analysis. Chimeras were removed using the function removeBimeraDenovo. The resulting chimera-checked, merged amplicon sequence variants (ASVs) were used for taxonomic classification using a BLAST against both the National Center for Biotechnology Information (NCBI) database and a custom BLAST database ^42^. The results were parsed into a table using the phyloseq package ^74^. Negative controls were assessed and the sum of reads from contaminating ASVs was subtracted from the samples. The number of reads identified as Kareniaceae species from each sample were calculated.

### MinION amplicon sequencing

The primers 18ScomF1 ^75^ and D2C ^71^ were used to amplify ∼3020 bp from select environmental samples as described in ^41^. This region comprising the almost complete sequence of the 18S through to the D2 region of the 28S rRNA gene. All PCR reactions were undertaken in 50 μL volumes composed of: 25 μL of MyFi™ Mix (Bioline), 2.5 μL of each primer (10 μM) and 5 μL of DNA extract. Cycling conditions were: 98 °C for 60 s, followed by 30 cycles of 98 °C for 10 s, 63 °C for 20 s, 72 °C for 90 s and a final extension of 72 °C for 10 min. Resulting amplicons were cleaned using Agencourt AMPure XP, and quantified by Qubit 3.0, using the 1X dsDNA high sensitivity kit (Thermo Fisher Scientific). The Oxford Nanopore Techologies (ONT) protocol for “Ligation sequencing amplicons - NativeBarcoding Kit 24 V14 (SQK-NBD114.24)” was performed following manufacturers’ instructions. The library was combined with loading beads and sequencing buffer before being transferred to a MinION FLO-MIN114 (R10.4.1) flow cell. The MinION was run for 36 h using the MinIT device. Negative control samples were prepared by using average sample volumes for each step to be representative of sample preparation. Local base calling of Fast5 files was performed using the MinIT (ONT-minit-release 19.05.2) device with the “flipflop” algorithm. Demultiplexing was performed using Porechop (v0.2.3). Analysis of the resulting sequences was performed on the custom alignment tool on the EPI2ME platform via the ONT website provided by Metrichor (Cambridge, United Kingdom).

FASTQ data was aligned to a custom reference list using Minimap2 (v2.28-r1209). The reference list comprised approximately 46 Kareniaceae sequences, each covering ∼700 bp of the D1 and D2 regions of the LSU rDNA. Alignments were filtered based on alignment score (AS: >400) and divergence (de: <0.05) using Bamtools, and a coverage report was generated with Samtools. Reads aligning to each species, as identified in the coverage report, were extracted and compiled into individual FASTQ files. The fastq files were submitted to NGSpeciesID (v0.2.1) to generate consensus sequences, with polishing performed using Medaka (v2.1.0). Consensus sequences were reviewed for orientation and aligned against the reference list using BioEdit (v7.2.5), followed by trimming.

### Phylogenetic Analysis

To verify identity of the large subunit RNA sequences we recovered from MinION amplicon sequencing and single-cell qPCR-amplified sequences, we inferred phylogenetic relationships of these sequences integrating high-quality LSU rDNA sequences from Kareniaceae taxa available in NCBI GenBank: 50 from diverse *Karenia* spp., and 12 from *Takayama* and *Karlodinium* taxa (6 per genus, as outgroup) (Supplementary Table 6). As a comparison to a prior HAB occurring in the same region ^57^, we re-analysed LSU rDNA sequences acquired during the 2014 event and inferred a phylogeny incorporating more taxa of *Takayama* and *Karlodinium* as outgroup (Supplementary Table 6).

For each sequence set, multiple sequence alignment was performed using MAFFT v7.471 ^76^ (*linsi* mode) at default setting. Spurious and non-phylogenetically informative sites in the alignment were removed using trimAl v1.5.0 ^77^ with the *-automated1* option. The trimmed alignment was used for phylogenetic inference using both maximum likelihood and Bayesian methods. Maximum likelihood tree was inferred using IQ-TREE v3.0.1 ^78^ with ultrafast bootstrap of 10,000 replicates (*-B 10000*), and incorporating ModelFinder to select the best-fit substitution model by default. Bayesian tree was inferred using MrBayes v3.2.7 ^79^ using the 4by4 nucleotide substitution model with gamma distribution (in four discrete categories), ran in four Markov chains over 2,500,000 generations (sample frequency 500 at default), and burn-in at 2000 samples.

### Brevetoxin analysis of water and culture samples using LC-MS/MS

Seawater samples (40 mL) were acidified by the addition of 10 µL of phosphoric acid (H□PO□). Subsequently, 5 mL of methyl tert-butyl ether (MTBE) was added, and the mixture was vigorously shaken to ensure thorough phase interaction. The samples were then centrifuged at 4000 × g for 5 minutes. Following centrifugation, the upper organic (MTBE) layer was carefully transferred to a 20 mL scintillation vial, avoiding carryover of the aqueous phase. An additional 1 mL of MTBE was gently added to the remaining sample and then removed and combined with the initial 5 mL extract. The combined MTBE extract was evaporated under a stream of nitrogen gas at 40□°C. The resulting residue was redissolved in 200 µL of methanol (MeOH) and transferred to an autosampler vial equipped with a 300 µL glass insert for subsequent analysis. he same protocol was followed for the sample of culture SM2, albeit with reduced volumes. Here 1.75 mL of culture was extracted with 1 mL MTBE, with the final extract resuspended in 200 µL MeOH and transferred into a glass insert for analysis.

LC-MS/MS analysis was performed using a Waters Xevo-TQS UPLC-MS instrument. For chromatographic separation a Waters Acquity BEH C8, 1.7 µm, 50 × 2.1 mm UPLC column held at 50 °C was used. The following gradient was used with a constant 0.3 mL min^-1^ flow rate (A: 0.1% ammonium hydroxide in water, B: MeOH). Initial conditions were 90% A (10% B) held for 0.5 min, with a linear change to 55% A at 1.5 min, followed a linear gradient to 10% A at 5 min. The solvent composition was then changed to 5% A at 5.25 min and held at this composition until 6.25 min. The column was then re-equilibrated using a linear gradient back to 90% A at 6.5 min, and then held until 8 min. The injection volume was 1 µL with samples held at 4 °C prior to injection. Positive electrospray ionization was used with the following optimised source parameters; capillary 3 kV, cone 50 V, source offset 50 V, desolvation temp 500 °C, desolvation gas 1200 L/hr, cone gas 150 L/hr, nebuliser gas 7 Bar, source temperature 150 °C. Quantitative multiple reaction monitoring (MRM) transitions, *m/z* precursor ion>product ion (collision energy, eV), for each toxin are shown in bold, with additional confirmation transitions also shown for each analyte monitored. Transitions are included for brevetoxins reported from microalgae and some B-type backbone shellfish metabolites: **BTX-1, 867.5>849.5 (12)**, 867.5>221.3 (20), 867.5>385.3 (20); **BTX-2, 895.5>301.2 (25)**, 895.5>319.2 (21), 895.5>859.4 (20), 895.5>877.4 (15); **BTX-3, 897.5>725.3 (18)**, 897.5>769.2 (15), 897.5>807.3 (10); **BTX-B5, 911.5>875.5 (19)**, 911.5>319.2 (32), 911.5>455.3 (26); **BTX-B1, 1018.5>248.1 (40)**, 1018.5>204.1 (52), 1018.5>222.1 (36), 1018.5>276.2 (41); **dBTX-B2, 1018.6>204.1 (44)**, 1018.6>248.2 (39), 1018.6>753.5 (30), 1018.6>929.5 (30); **BTX-B2, 1034.6>929.5 (31)**, 1034.6>911.3 (32), 1034.6>947.3 (33); **BTX-B4 (C16 fatty acid), 1272.7>326.3 (43)**, 1272.7>239.3 (46), 1272.7>929.7 (37), 1272.7>1016.5 (38). For instrument calibration, working standards containing various brevetoxin analogues were prepared as a 5-point calibration series from 2 ng/mL to 50 ng/mL, which was determined to be linear over this range. Samples with brevetoxin levels above the calibration range were diluted and rerun, or a higher concentration standard prepared if the calibration is still linear. For those analogues that did not have calibration material, they were calibrated of the nearest structurally related analogue (Supplementary Table 7).

### Rotifer and brine shrimp mortality assays

Whole-organism toxicity was assessed using acute mortality assays with rotifers (*Brachionus plicatilis*; 24 h) and brine shrimp (*Artemia salina* 2-3 day old nauplii; 72 h). Assays were conducted concurrently using a non-clonal, stationary-phase culture of *Karenia cristata* (SM2) and a field-collected bloom sample from West Lakes (Fig. 1a, site 24) collected on 14 January 2026, when high cell densities were persisting. Prior to exposure, algal cell concentrations were determined from 1 mL subsamples from each algal stock (15.6 × 10□cells L□¹ for SM2 and 3.02 × 10□cells L□¹ for the West Lakes sample). Exposures were conducted in 24-well plates using 1:3 serial dilutions of whole-cell algal suspensions prepared in artificial seawater (salinity 36). Each dilution series was replicated four times. Each well contained 2.0 mL of test solution, to which 0.4 mL of a rotifer or Artemia stock suspension was added to initiate exposure, yielding on average 12 rotifers or 8 Artemia nauplii per well. Rotifer assays were conducted under ambient laboratory temperature (∼21 °C) and lighting, whereas the longer Artemia assays were incubated under controlled conditions (12:12 h light:dark cycle, 17 °C). Control wells contained artificial seawater only. Algal cell concentrations in Artemia assays were verified at the end of the exposure period to confirm sustained exposure. Concentration–response relationships were modelled in R using three-parameter log-logistic functions implemented in the drm() function of the drc package, with LC10 and LC50 values estimated from fitted models

### RTgill-W1 viability assay

The toxicity of the supernatant of the same non-clonal culture of *Karenia cristata* (SM2) was assessed against RTgill-W1 cells. The algae was grown at 17□C in a 500ml Erlenmeyer flask containing 250ml of K media at a salinity of 35 under 12:12 light:dark conditions. Once stationary phase had been reached, the culture was gently mixed, a one millilitre subsample was fixed with lugols and counted on a Sedgewick rafter to determine the algal density, Another 50ml sub sample was centrifuged in the dark at 4□C at 1000g for 5 mins to acquire a cell-free supernatant. The supernatant was diluted in step-wise 2X dilutions in K media before diluting all concentrations another 4X in L15/ex, resulting in RTgill-W1 exposure concentrations ranging from 16X the algal stock culture concentration down to 2048X of that level, in 25% K media.

The *Oncorhynchus mykiss* rainbow trout cell line (RTgill-W1, CRL-2523, ATCC, USA), was cultured following ATTC recommendation and the method of 47 and 48. Bioassays were performed and analysed according to 47 and 48. Statistical and graphical analyses for the bioassays were conducted in R version 4.4.080. To examine the relationship between viability and microalgal cell concentration, nonlinear dose–response models were fitted using the drm function from the drc package in RStudio 2026.01.0 Build 392. Multiple candidate models were compared using Akaike’s Information Criterion (AIC), and the model with the lowest AIC was selected as the best fit. Parameter estimates and goodness-of-fit metrics were extracted from the selected LL3 model.

## Acknowledgements

We thank Sam Gaylard (SA EPA), Phil Dunne, Anke-Maria Hoeffer, Carolyn Ricci, Tracey Spokes, Serhii Snihirov and Nicola Lieff for sample collection. Jacob Thomson-Laing, Karthiga Kumanan, John Pearman (Cawthron Institute) and Rosie Buselli (UTS) helped with molecular analyses. Sandrin Feig and Olivier Bibari from the University of Tasmania and Maria Byrne from the University of Sydney provided help in sample fixation and Scanning Electron Microscopy. Lucy Thompson (Cawthron Institute) assisted with culturing and microscopy. Grant Pitcher (South Africa) and Nicolas Chomerat (France) provided helpful information. We thank the captain and crew of the RV Ngerin and FPV Southern Ranger. Satellite data was sourced from Australia’s Integrated Marine Observing System (IMOS), enabled by the National Collaborative Research Infrastructure Strategy (NCRIS). It is operated by a consortium of institutions as an unincorporated joint venture, with the University of Tasmania as Lead Agent.

