## Supplementary material for "A catastrophic marine mortality event caused by a complex algal bloom including the novel brevetoxin producer, *Karenia cristata* (Dinophyceae)"

**Supplementary information**

Supplementary Table 1. *Karenia* blooms involved in marine mass mortalities (updated after Brand et al. 2012)

| **Species** | **Species first described** | **Fish and invertebrates impacted** | **Seabirds impacted** | **Mammal mortalities** | **Human impacts** | **Toxins** | **Distribution** | **Reference** |
| --- | --- | --- | --- | --- | --- | --- | --- | --- |
| *K. brevis* | Florida, USA (Davis 1948) | crabs, fish, invertebrates, scallops, shrimp, turtles | lesser scaup, cormorants, mergansers | manatees, dolphins, dogs, coyotes | NSP, respiratory distress | Brevetoxins  Brevisamide  Brevisin  Brevenal | Gulf of Mexico (since 1500s) | Steidinger et al.1998 |
| *K. brevisulcata* | New Zealand ([Chang 1999)](https://www.sciencedirect.com/science/article/pii/S156898831100148X#bib0210) | abalone, clams, marlin, mussels, swordfish, sea urchins, tuna | no | no | respiratory distress, eye and skin irritation | Brevisulcenals  Brevisulcatic acids | New Zealand 1988 | Chang 1999 |
| *K. concordia* | New Zealand ([Chang and Ryan 2004)](https://www.sciencedirect.com/science/article/pii/S156898831100148X#bib0220) | abalone, eel, goby, flounder, mullet, parore, spotty | no | no | NSP-like symptoms | not characterized | New Zealand | Chang & Ryan 2004 |
| *K. cristata* | South Africa ([Botes et al. 2003)](https://www.sciencedirect.com/science/article/pii/S156898831100148X#bib0090) | abalone, chitons, limpets, mussels, octopus, periwinkle, whelks, sardine, starfish | no | no | respiratory distress, eye and skin irritation | not characterized | South Africa 1988, 1989, 1995-96 | Horstman et al.1991; Pitcher & Matthews 1996 |
| *K. mikimotoi* | Japan ([Oda 1935)](https://www.sciencedirect.com/science/article/pii/S156898831100148X#bib1070) | abalone, bivalves, cockles, crabs, echinoderms, eel, flat fish, holothurians, lobster, lugworm, mussels, salmon, sea urchins, scallops, starfish, sting rays, turbo shells | no | no | none known | Gymnocin-A  Gymnocin-B  PUFA ? | Australia 1995, 2014, Europe (Ireland, Norway, Scotland), China, Japan, Hong Kong, Korea, North America | Davidson et al. 2009, PIRSA 2014 |
| *K. selliformis* | New Zealand ([Haywood et al. 2004)](https://www.sciencedirect.com/science/article/pii/S156898831100148X#bib0565) | ascidians, bivalves, chiton, crustaceans, flounder, gastropods, jellyfish, octopus, oysters, salmon, sea anemones, sea cucumber, sea urchins, sponges, starfish, surf clams | no | none known | none known | Gymnodimine (not all strains) | New Zealand 1993-96, Chile 1999, China 2023, Japan 2021, Kuwait 1999, Russia, Tunesia | Iwataki et al. 2022, Mardones et al. 2020, Orlova et al. 2022, Xu et al. 2025 |
| *K. umbella* | Tasmania (de Salas et al 2004) | Rainbow trout, salmon | no | none known | none known | not characterized | Tasmania, New Zealand | De Salas et al 2004  Rolton et al 2022 |
| ***K.cristata***  *K. mikimotoi* |  | Many documented^2^. Some of these include : abalone, boarfish, clams, cockles, crabs, cobbler, cowfish, eels, flathead, globe fish, octopus, leatherjacket, lobster, morwong, mussels, pufferfish, sharks, silverbelly, rays, salmon, scallops, sea cucumbers, sea dragons, sea urchins, snails, sponges, toadfish, whiting, worms | Not yet confirmed. Several suspected | Not yet confirmed, several suspected, including kangaroos | respiratory irritation, eye & skin irritation, cough, | Brevetoxins  Other toxins not yet confirmed | South Australia 2025 | **Present work** |

Supplementary Table 2. Sites and dates of marine water sample collection from South Australia, March – August 2025. (SA EPA = South Australian Environmental Protection Authority, SARDI = South Australian Research and Development Institute, PIRSA = Department of Primary Industries and Regions in South Australia, i = Illumina metabarcoding, m = MinION analysis, q = qPCR analysis, * = sample collected on filters and preserved in Longmire’s buffer, ^ = samples preserved with Lugol's Iodine, DCM = deep chlorophyll maximum)

| **Site number** | **Location** | **Sampler** | **Depth** | **Date** | **Latitude (°S)** | **Longitude (°E)** |
| --- | --- | --- | --- | --- | --- | --- |
| 1 | Encounter Bay Boat Ramp^i,m^ | SA EPA/SARDI | Surface | 18/03/2025^q^ | 35.585 | 138.598 |
| 2 | Petrel Cove Beach^i,m^ | SA EPA/SARDI | Surface | 18/03/2025^q^ | 35.593 | 138.600 |
| 3 | Waitpinga Beach^i,m^ | SA EPA/SARDI | Surface | 18/03/2025^q^ | 35.635 | 138.499 |
| 4 | Parsons Headland^i,m^ | SA EPA/SARDI | Surface | 18/03/2025^q^ | 35.634 | 138.475 |
| 5 | Goolwa Beach Site 3^i^ | PIRSA, Clinton Wilkinson | Surface | 7/05/2025^q^ | 35.539 | 138.819 |
| 6 | Goolwa Beach Site 4^i^ | PIRSA, Clinton Wilkinson | Surface | 7/05/2025^q^ | 35.551 | 138.851 |
| 7 | Stokes Bay, Kangaroo Island^i,m^ | PIRSA, Clinton Wilkinson | Surface | 13/05/2025^q^ | 35.625 | 137.159 |
| 8 | Point Boston^^,m^ | PIRSA, Clinton Wilkinson | Surface | 6/06/2025^q^ | 34.649 | 135.935 |
| 9 | Emu Bay, Kangaroo Island^^,m^ | PIRSA, Clinton Wilkinson | Surface | 13/06/2025^q^ | 35.590 | 137.506 |
| 10 | Emu Bay, Kangaroo Island^^,m^ | PIRSA, Clinton Wilkinson | Surface | 13/06/2025^q^ | 35.590 | 137.506 |
| 11 | SARDI S/W Intakes^^,m^ | SARDI | 1m | 27/06/2025^q^ | 34.958 | 138.489 |
| 12 | Ardrossan Jetty^^,m^ | PIRSA, Clinton Wilkinson | Surface | 9/07/2025^q^ | 34.425 | 137.925 |
| 13 | Port River Dock | SA EPA | Surface | 14/07/2025^q^ | 34.761 | 138.510 |
| 14 | West Lakes Inlet | SA EPA | Surface | 14/07/2025^q^ | 34.878 | 138.486 |
| 15 | Franklin Harbour | PIRSA, Clinton Wilkinson | Surface | 28/07/2025^q^ | 33.752 | 136.905 |
| 16 | Stansbury (slick on water)^^,m^ | PIRSA, Clinton Wilkinson | Integrated tube (4m) | 7/07/2025^q^ | 34.889 | 137.840 |
| 17 | Stansbury^ | PIRSA, Clinton Wilkinson | Integrated tube (4m) | 12/05/2025^m,q^ | 34.898 | 137.826 |
|  |  |  |  | 28/07/2025^q^ |  |  |
| 18 | O'Sullivan Beach Boat Ramp* | Nikola Streiber, Phil Dunne, Anke-Maria Hoeffer, Neil MacDonald, C Ricii, T Spokes, Shauna Murray | Surface | 27/07/2025^q^ | 35.120 | 138.467 |
|  |  |  |  | 3/08/2025^q^ |  |  |
|  |  |  |  | 10/08/2025^q^ |  |  |
|  |  |  |  | 17/08/2025^q^ |  |  |
|  |  |  |  | 24/08/2025^q^ |  |  |
|  |  |  |  | 31/08/2025^q^ |  |  |
|  |  |  |  | 6/09/2025^q^ |  |  |
|  |  |  |  | 13/09/2025^q^ |  |  |
| 19 | Port Noarlunga Jetty* | Nikola Streiber, Phil Dunne, Anke-Maria Hoeffer, Neil MacDonald, C Ricii, T Spokes, Shauna Murray | Surface | 27/07/2025^q^ | 35.149 | 138.468 |
|  |  |  |  | 3/08/2025^q^ |  |  |
|  |  |  |  | 10/08/2025^q^ |  |  |
|  |  |  |  | 17/08/2025^q^ |  |  |
|  |  |  |  | 24/08/2025^q^ |  |  |
|  |  |  |  | 31/08/2025^q^ |  |  |
|  |  |  |  | 6/09/2025^q^ |  |  |
|  |  |  |  | 13/09/2025^q^ |  |  |
| 20 | Seaford Beach* | Nikola Streiber, Phil Dunne, Anke-Maria Hoeffer, Neil MacDonald, C Ricii, T Spokes | Surface | 27/07/2025^q^ | 35.181 | 138.468 |
|  |  |  |  | 3/08/2025^q^ |  |  |
|  |  |  |  | 10/08/2025^q^ |  |  |
|  |  |  |  | 17/08/2025^q^ |  |  |
|  |  |  |  | 24/08/2025^q^ |  |  |
|  |  |  |  | 31/08/2025^q^ |  |  |
|  |  |  |  | 6/09/2025^q^ |  |  |
| 21 | Moana Beach* | Nicola Lieff | Surface | 31/07/2025 | 35.198 | 138.470 |
|  |  |  |  | 3/08/2025 |  |  |
|  |  |  |  | 19/08/2025 |  |  |
|  |  |  |  | 31/08/2025 |  |  |
| 22 | Maslin Beach* | Nicola Lieff | Surface | 3/08/2025 | 35.234 | 138.471 |
|  |  |  |  | 19/08/2025 |  |  |
|  |  |  |  | 31/08/2025 |  |  |
| 23 | Aldinga beach* | Nicola Lieff | Surface | 3/08/2025 | 35.290 | 138.444 |
|  |  |  |  | 19/08/2025 |  |  |
|  |  |  |  | 31/08/2025 |  |  |
| 24 | West Lakes* | Anastasiia Snigirova, Serhii Snihirov, Shauna Murray | Surface | 3/08/2025^q^ | 34.875 | 138.485 |
|  |  |  |  | 6/09/2025 |  |  |
| 25 | Port River* | Anastasiia Snigirova, Serhii Snihirov, Shauna Murray | Surface | 3/08/2025^q^ | 34.770 | 138.514 |
|  |  |  |  | 5/09/2025 |  |  |
| 26 | St Vincent Gulf^i,m^ | SARDI –RV Ngerin | DCM | 22/05/2025^q^ | 34.786 | 138.300 |
| 27 | St Vincent Gulf^i,m^ | SARDI –RV Ngerin | DCM | 22/05/2025^q^ | 35.184 | 137.614 |
| 28 | St Vincent Gulf^i,m^ | SARDI –RV Ngerin | DCM | 22/05/2025^q^ | 34.940 | 137.880 |
| 29 | St Vincent Gulf^i,m^ | SARDI –RV Ngerin | DCM | 22/05/2025^q^ | 35.137 | 137.878 |
| 30 | St Vincent Gulf^i,m^ | SARDI –RV Ngerin | DCM | 23/05/2025^q^ | 34.786 | 138.150 |
| 31 | St Vincent Gulf^i,m^ | SARDI –RV Ngerin | DCM | 23/05/2025^q^ | 34.786 | 137.980 |
| 32 | St Vincent Gulf | PIRSA FPV Southern Ranger | 1m | 16/07/2025^q^ | 34.531 | 138.049 |
| 33 | St Vincent Gulf | PIRSA FPV Southern Ranger | 1m | 16/07/2025 | 34.630 | 138.113 |
| 34 | St Vincent Gulf | PIRSA FPV Southern Ranger | 1m | 16/07/2025^q^ | 34.674 | 138.358 |
| 35 | St Vincent Gulf | PIRSA FPV Southern Ranger | 1m | 16/07/2025^q^ | 34.749 | 138.361 |
| 36 | St Vincent Gulf | PIRSA FPV Southern Ranger | 1m | 16/07/2025^q^ | 34.880 | 138.122 |
| 37 | St Vincent Gulf | PIRSA FPV Southern Ranger | 1m | 16/07/2025^q^ | 35.249 | 138.441 |
| 38 | St Vincent Gulf | PIRSA FPV Southern Ranger | 1m | 16/07/2025^q^ | 35.351 | 138.353 |
| 39 | St Vincent Gulf | PIRSA FPV Southern Ranger | 1m | 16/07/2025^q^ | 35.457 | 138.142 |

Supplementary Table 3. Strains of *Karenia* species and *Karlodinium veneficum* in culture used as standards for comparison in the qPCR assays. (The CAW code refers to isolates from the Cawthron Institute Culture Collection of Microalgae)

| **Strain number** | **Species name** |
| --- | --- |
| CAWD80 | *Karenia bidigitata* |
| CAWD81 | *Karenia bidigitata* |
| CAWD92 | *Karenia bidigitata* |
| CAWD04 | *Karenia brevis* |
| CAWD6 | *Karenia brevis* |
| CAWD08 | *Karenia brevis* |
| CAWD122 | *Karenia brevis* |
| CAWD82 | *Karenia brevisulcata* |
| CAWD457 | *Karenia longicanalis* |
| CAWD05 | *Karenia mikimotoi* |
| CAWD63 | *Karenia mikimotoi* |
| CAWD117 | *Karenia mikimotoi* |
| CAWD133 | *Karenia mikimotoi* |
| CAWD134 | *Karenia mikimotoi* |
| CAWD192 | *Karenia mikimotoi* |
| CAWD91 | *Karenia papilionacea* |
| CAWD79 | *Karenia selliformis* |
| CAWD93 | *Karlodinium veneficum* |
| CAWD84 | *Karlodinium veneficum* |

Supplementary Table 4. Summary of Illumina metabarcoding data. Number of sequence reads identified as Kareniaceae species from environmental samples collected from South Australia.

| **Site number** | **Location** | **Date** | ***Karenia cristata*** | ***Karenia mikimotoi*** | ***Karenia longicanalis*** | ***Karlodinium* sp. 1** | ***Karlodinium* sp. 2** | ***Karlodinium veneficum*** |
| --- | --- | --- | --- | --- | --- | --- | --- | --- |
| 1 | Encounter Bay Boat Ramp | 18/03/2025 | 865 | 2925 | 56 | 0 | 0 | 0 |
| 2 | Petrel Cove Beach | 18/03/2025 | 15679 | 1023 | 0 | 0 | 168 | 0 |
| 3 | Waitpinga Beach | 18/03/2025 | 14398 | 1347 | 0 | 252 | 137 | 0 |
| 4 | Parsons Headland | 18/03/2025 | 1520 | 689 | 0 | 163 | 41 | 0 |
| 5 | Goolwa Beach Site 3 | 7/05/2025 | 0 | 0 | 0 | 36 | 0 | 0 |
| 6 | Goolwa Beach Site 4 | 7/05/2025 | 1158 | 0 | 0 | 115 | 0 | 0 |
| 7 | Stokes Bay, Kangaroo Island | 13/05/2025 | 739 | 1028 | 15 | 137 | 0 | 23 |
| 26 | St Vincent Gulf | 22/05/2025 | 91874 | 858 | 0 | 85 | 0 | 0 |
| 27 | St Vincent Gulf | 22/05/2025 | 0 | 0 | 0 | 17 | 0 | 0 |
| 28 | St Vincent Gulf | 22/05/2025 | 121236 | 0 | 0 | 0 | 0 | 8 |
| 29 | St Vincent Gulf | 22/05/2025 | 255 | 0 | 0 | 181 | 0 | 0 |
| 30 | St Vincent Gulf | 23/05/2025 | 47 | 15164 | 232 | 0 | 154 | 0 |
| 31 | St Vincent Gulf | 23/05/2025 | 558 | 400 | 0 | 26 | 0 | 0 |

Supplementary Table 5. Primers and probes used in the qPCR assays.

| **Species** | **Primer and probe name and sequence** | **Final Concentration** | **Reference** |
| --- | --- | --- | --- |
| *Karenia brevisulcata* | KBS460-F: GATCTGGATGCGATACTGAAT | 300 nM | Smith et al 2014 |
|  | KBS585-R: AGCACTGCTACAAGACATATAA | 900 nM |  |
|  | KBS544-P: 6-FAM/TG ACT GAA T/ZEN/G TCC CTA GTT GAA CTC /IABkFQ/ | 100 nM |  |
| *Karenia mikimotoi* | KM541-F: CGAGTGACTGAATGTCCTCA | 500 nM | Smith et al 2014 |
|  | KM645-R: CCAACAACCTTCATGCAGAG | 250 nM |  |
|  | KM578-P: 6-FAM/CT ACC AGA C/ZEN/A CAC AGA GAG CAG /IABkFQ | 100 nM |  |
| *Karenia papilionacea* | KP449-F: TCTGGATGCGATACTGGTTG | 1000 nM | Smith et al 2014 |
|  | KP682-R: TACTTATGTCAAGGATGTGTTC | 750 nM |  |
|  | KP630-P: 6-FAM/CT TGT TAG T/ZEN/T ACC TGG CAT GAG AC/IABkFQ | 125 nM |  |
| *Karenia longicanalis /umbella* | KU480-F: ATGTCAACGTCAGTTCACAAT | 750 nM | Smith et al 2014 |
|  | KU623-R: GCACGAGACGAGGCTTA | 250 nM |  |
|  | KU542-P: 6-FAM/TT CGA CTA G/ZEN/G CAC ATT CAG TCA C/IABkFQ | 100 nM |  |
| *Karenia brevis* | Kb-rbcl-F: TGAAACGTTATTGGGTCTGT | 400 nM | Gray et al 2003* |
|  | Kb-rbcl-R: AGGTACACACTTTCGTAAACTA | 400 nM |  |
|  | Kb-rbcl-P: 6-FAM/AC GAA TTA A/ZEN/C CTT AGT CTC GGG TTA /IABkFQ | 200 nM |  |

*Probe sequence was edited from the published paper to be adapted for a FAM TaqMan format.

Supplementary Table 6. Kareniaceae reference LSU rDNA sequences used for phylogenetic analyses in this study.

| **NCBI accession** | **Species** | **Strain/isolate** | **Sequence name used in the analysis** | **Source location** | **Inclusion for analysis** |
| --- | --- | --- | --- | --- | --- |
| AY590123.1 | *Karenia asterichroma* |  | Karenia_asterichroma_AY590123 | Australia: Tasmania, Pirates Bay | All trees |
| AY947662.1 | *Karenia bidigitata* | CAWD80 | Karenia_bidigitata_CAWD80_AY947662 | New Zealand: Foveaux Strait | All trees |
| AY947663.1 | *Karenia bidigitata* | CAWD81 | Karenia_bidigitata_CAWD81_AY947663 | New Zealand: Foveaux Strait | All trees |
| EU165308.1 | *Karenia brevis* | CCMP2228 | Karenia_brevis_CCMP2228_EU165308 | NA | All trees |
| MW177918.1 | *Karenia brevis* | CAWD122 | Karenia_brevis_CAWD122_MW177918 | USA: Pensacola Beach, Florida | All trees |
| AY243032.1 | *Karenia brevisulcata* |  | Karenia_brevisulcata_AY243032 | New Zealand: Cawthron Institute | All trees |
| KJ508359.1 | *Karenia brevisulcata* | IFR1133 | Karenia_brevisulcata_IFR1133_KJ508359 | France: Concarneau Bay | All trees |
| AY243963.1 | *Karenia cristata* |  | Karenia_cristata_AY243963 | NA | All trees |
| KJ508360.1 | *Karenia cristata* | IFR13-067 | Karenia_cristata_IFR13-067_KJ508360 | Saint Pierre and Miquelon | All trees |
| OR527549.1 | *Karenia hui* | RCHA6401-H7-LSU | Karenia_hui_RCHA6401-H7-LSU_OR527549 | China | All trees |
| OR527550.1 | *Karenia hui* | RCHA6402-C11-LSU | Karenia_hui_RCHA6402-C11-LSU_OR527550 | China | All trees |
| KY287670.1 | *Karenia longicanalis* | HK01 | Karenia_longicanalis_HK01_KY287670 | China | All trees |
| LC671812.1 | *Karenia longicanalis* | LAkKL297 | Karenia_longicanalis_LAkKL297_LC671812 | Japan | All trees |
| LC671814.1 | *Karenia longicanalis* | LAkKL299 | Karenia_longicanalis_LAkKL299_LC671814 | Japan | All trees |
| LC671815.1 | *Karenia longicanalis* | LAkKL300 | Karenia_longicanalis_LAkKL300_LC671815 | Japan | All trees |
| MG737367.1 | *Karenia longicanalis* | GSLJ | Karenia_longicanalis_GSLJ_MG737367 | Fuzhou, East China Sea | All trees |
| MZ465594.1 | *Karenia longicanalis* | K.lon_5 | Karenia_longicanalis_K.lon_5_MZ465594 | Russia: Khalaktyrsky beach, Avachinsky bay | All trees |
| AY263962.1 | *Karenia longicanalis (Karenia umbella)* | KULV01 | Karenia_longicanalis_umbella_KULV01_AY263962 | Australia: Spring Bay, Tasmania | All trees |
| AY263963.1 | *Karenia longicanalis (Karenia umbella)* | KUTN05 | Karenia_longicanalis_umbella_KUTN05_AY263963 | Australia: Taranna, Tasmania | All trees |
| AY266329.1 | *Karenia longicanalis (Karenia umbella)* | GY2DE | Karenia_longicanalis_umbella_GY2DE_AY266329 | Australia: River Derwent, Tasmania | All trees |
| KJ508372.1 | *Karenia longicanalis (Karenia umbella)* | IFR13-377 | Karenia_longicanalis_umbella_IFR13-377_KJ508372 | Saint Pierre and Miquelon | All trees |
| EF469238.1 | *Karenia mikimotoi* | KMWL01 | Karenia_mikimotoi_KMWL01_EF469238 | Australia: West Lakes, South Australia | All trees |
| KJ508361.1 | *Karenia mikimotoi* | IFR980 | Karenia_mikimotoi_IFR980_KJ508361 | France: Gulf of Lions | All trees |
| KJ508365.1 | *Karenia mikimotoi* | IFR11-056 | Karenia_mikimotoi_IFR11-056_KJ508365 | New Caledonia | All trees |
| MT754546.1 | *Karenia mikimotoi* | HK-17 | Karenia_mikimotoi_HK-17_MT754546 | NA | All trees |
| OQ781172.1 | *Karenia mikimotoi* | voucher17-K.mikimotoi | Karenia_mikimotoi_voucher17-K.mikimotoi_OQ781172 | NA | All trees |
| OR527546.1 | *Karenia mikimotoi* | RCHA6001 | Karenia_mikimotoi_RCHA6001_OR527546 | China | All trees |
| OR527548.1 | *Karenia mikimotoi* | RCHA6002-17km-LSU | Karenia_mikimotoi_RCHA6002-17km-LSU_OR527548 | China | All trees |
| AB623227.1 | *Karenia papilionacea* | KP02URA | Karenia_papilionacea_KP02URA_AB623227 | Japan: Kouchi, Uranouchi Bay | All trees |
| AY590124.1 | *Karenia papilionacea* |  | Karenia_papilionacea_AY590124 | Australia: Tasmania, Moulting Bay | All trees |
| FN649411.1 | *Karenia papilionacea* | VGO679 | Karenia_papilionacea_VGO679_FN649411 | Spain | All trees |
| KJ508366.1 | *Karenia papilionacea* | IFR562 | Karenia_papilionacea_IFR562_KJ508366 | France: Douarnenez Bay | All trees |
| KJ508367.1 | *Karenia papilionacea* | IFR13-294 | Karenia_papilionacea_IFR13-294_KJ508367 | Saint Pierre and Miquelon | All trees |
| LC055217.1 | *Karenia papilionacea* | KpSIK12H | Karenia_papilionacea_KpSIK12H_LC055217 | Japan:Oita | All trees |
| LC055220.1 | *Karenia papilionacea* | KspNOM6H | Karenia_papilionacea_KspNOM6H_LC055220 | Japan:Kochi | All trees |
| MG737370.1 | *Karenia papilionacea* | KP01 | Karenia_papilionacea_KP01_MG737370 | Xiamen, East China Sea | All trees |
| OR527552.1 | *Karenia papilionacea* | RCHA6203-TLJ-H1-LSU | Karenia_papilionacea_RCHA6203-TLJ-H1-LSU_OR527552 | China | All trees |
| OR527553.1 | *Karenia papilionacea* | RCHA6202-DYW-K-LSU | Karenia_papilionacea_RCHA6202-DYW-K-LSU_OR527553 | China | All trees |
| PP951872.1 | *Karenia papilionacea* | KW-E1-14 | Karenia_papilionacea_KW-E1-14_PP951872 | Kuwait: Kuwait Bay | All trees |
| U92252.1 | *Karenia papilionacea* |  | Karenia_papilionacea_U92252 | NA | All trees |
| LC671820.1 | *Karenia selliformis* | MoKr600 | Karenia_selliformis_MoKr600_LC671820 | Japan | All trees |
| LC671821.1 | *Karenia selliformis* | KKsKs74 | Karenia_selliformis_KKsKs74_LC671821 | Japan | All trees |
| LC671827.1 | *Karenia selliformis* | M3 | Karenia_selliformis_M3_LC671827 | Japan | All trees |
| LC671830.1 | *Karenia selliformis* | M6 | Karenia_selliformis_M6_LC671830 | Japan | All trees |
| LC671833.1 | *Karenia selliformis* | LK2Ks306 | Karenia_selliformis_LK2Ks306_LC671833 | Japan | All trees |
| LC671836.1 | *Karenia selliformis* | AsahiL7 | Karenia_selliformis_AsahiL7_LC671836 | Japan | All trees |
| MN203221.1 | *Karenia selliformis* | CREAN_KS02 | Karenia_selliformis_CREAN_KS02_MN203221 | NA | All trees |
| MZ465598.1 | *Karenia selliformis* | K.sell_16 | Karenia_selliformis_K.sell_16_MZ465598 | Russia: Utashud Island | All trees |
| KJ508369.1 | *Karenia* sp. | IFR868 | Karenia_sp_IFR868_KJ508369 | France: Leucate lagoon | All trees |
| KJ508373.1 | *Karenia* sp. | IFR528 | Karenia_sp_IFR528_KJ508373 | France: Douarnenez Bay | All trees |
| EF469234.1 | *Karlodinium antarcticum* | KDANSO10 | Karlodinium_antarcticum_KDANSO10_EF469234 | Australia: the Southern Ocean E146 S51 | All trees |
| DQ114467.1 | *Karlodinium armiger* |  | Karlodinium_armiger_DQ114467 | NA | Complete HAB tree only |
| KJ508375.1 | *Karlodinium armiger* | IFR-KAR-01D | Karlodinium_armiger_IFR-KAR-01D_KJ508375 | France: Diana lagoon | Complete HAB tree only |
| DQ151560.1 | *Karlodinium australe* | KDATL11 | Karlodinium_australe_KDATL11_DQ151560 | Australia: N.S.W., Tuggerah Lakes | All trees |
| LC521279.1 | *Karlodinium australe* | GBOKZ88 | Karlodinium_australe_GBOKZ88_LC521279 | Japan: Mikawa Bay | Complete HAB tree only |
| OR842796.1 | *Karlodinium australe* | BGERL277 | Karlodinium_australe_BGERL277_OR842796 | China: Fangcheng | Complete HAB tree only |
| LC521281.1 | *Karlodinium azanzae* | GBMB68 | Karlodinium_azanzae_GBMB68_LC521281 | Philippines: Manila Bay | Complete HAB tree only |
| EF469232.1 | *Karlodinium ballantinum* | KDBMP01 | Karlodinium_ballantinum_KDBMP01_EF469232 | Australia: Mercury Passage, Tasmania | Complete HAB tree only |
| PV240841.1 | *Karlodinium ballantinum* | UNC1840 | Karlodinium_ballantinum_UNC1840_PV240841 | Pacific Ocean: Station Papa | All trees |
| EF469231.1 | *Karlodinium conicum* | KDCSO15 | Karlodinium_conicum_KDCSO15_EF469231 | Australia: the Southern Ocean E147 S45 | All trees |
| EF469233.1 | *Karlodinium corrugatum* | KDGSO08 | Karlodinium_corrugatum_KDGSO08_EF469233 | Australia: the Southern Ocean E145 S53 | Complete HAB tree only |
| EF469235.1 | *Karlodinium decipiens* | KDDSO10 | Karlodinium_decipiens_KDDSO10_EF469235 | Australia: the Southern Ocean E146 S51 | Complete HAB tree only |
| LC521288.1 | *Karlodinium decipiens* | MD482 | Karlodinium_decipiens_MD482_LC521288 | Japan: Sagami Bay | Complete HAB tree only |
| MZ358884.1 | *Karlodinium digitatum* | KdLomme04 | Karlodinium_digitatum_KdLomme04_MZ358884 | NA | Complete HAB tree only |
| MZ358887.1 | *Karlodinium digitatum* | KdLomme01 | Karlodinium_digitatum_KdLomme01_MZ358887 | NA | Complete HAB tree only |
| MT161377.1 | *Karlodinium elegans* | B601 | Karlodinium_elegans_B601_MT161377 | China | All trees |
| KJ508379.1 | *Karlodinium gentienii* | IFR12-234 | Karlodinium_gentienii_IFR12-234_KJ508379 | France: Concarneau Bay | All trees |
| LC521290.1 | *Karlodinium gentienii* | LUR99 | Karlodinium_gentienii_LUR99_LC521290 | Japan: Okinawa | Complete HAB tree only |
| LC521291.1 | *Karlodinium gentienii* | GBJS47 | Karlodinium_gentienii_GBJS47_LC521291 | Japan: Miura | Complete HAB tree only |
| DQ114466.1 | *Karlodinium veneficum* |  | Karlodinium_veneficum_DQ114466 | NA | Complete HAB tree only |
| KJ508380.1 | *Karlodinium veneficum* | IFR10-150 | Karlodinium_veneficum_IFR10-150_KJ508380 | France: Dinan Kerloc'h Bay | All trees |
| KJ508381.1 | *Karlodinium veneficum* | IFR-KVE-01D | Karlodinium_veneficum_IFR-KVE-01D_KJ508381 | France: Diana lagoon | Complete HAB tree only |
| PP565094.1 | *Karlodinium veneficum* | VGO1111 | Karlodinium_veneficum_VGO1111_PP565094 | NA | Complete HAB tree only |
| MG737358.1 | *Karlodinium zhouanum* | TIO397 | Karlodinium_zhouanum_TIO397_MG737358 | Daya bay, South China Sea | Complete HAB tree only |
| MZ358878.1 | *Karlodinium zhouanum* | KzLomme01 | Karlodinium_zhouanum_KzLomme01_MZ358878 | NA | Complete HAB tree only |
| DQ656116.1 | *Takayama acrotrocha* | GT15 | Takayama_acrotrocha_GT15_DQ656116 | Singapore | Complete HAB tree only |
| FJ024702.1 | *Takayama acrotrocha* | MC728-B4 | Takayama_acrotrocha_MC728-B4_FJ024702 | Italy: Gulf of Naples | Complete HAB tree only |
| OR778358.1 | *Takayama acrotrocha* | BGERL274 | Takayama_acrotrocha_BGERL274_OR778358 | China: Beihai | All trees |
| AY284951.1 | *Takayama helix* | TTPA01 | Takayama_helix_TTPA01_AY284951 | Australia: Port Arthur, TAS | All trees |
| LC535355.1 | *Takayama helix* | ThK1T10 | Takayama_helix_ThK1T10_LC535355 | Japan:Nagasaki, Off-Mie | Complete HAB tree only |
| MZ358882.1 | *Takayama helix* | ThLomme01 | Takayama_helix_ThLomme01_MZ358882 | NA | Complete HAB tree only |
| KJ508388.1 | *Takayama* sp. | IFR909 | Takayama_sp_IFR909_KJ508388 | France: Quiberon Bay | All trees |
| AY284948.1 | *Takayama tasmanica* | TTDE01 | Takayama_tasmanica_TTDE01_AY284948 | Australia: River Derwent, TAS | All trees |
| MZ358879.1 | *Takayama tasmanica* | TtLomme02 | Takayama_tasmanica_TtLomme02_MZ358879 | NA | Complete HAB tree only |
| MZ358880.1 | *Takayama tasmanica* | TtLomme01 | Takayama_tasmanica_TtLomme01_MZ358880 | NA | Complete HAB tree only |
| EF469230.1 | *Takayama tuberculata* | TTBSO11.1 | Takayama_tuberculata_TTBSO11_EF469230 | Australia: the Southern Ocean E146 S49 | All trees |
| AY764178.1 | *Takayama xiamenensis* | TPXM | Takayama_xiamenensis_TPXM_AY764178 | NA | All trees |

Supplementary Table 7. Brevetoxin calibration standards used in toxin testing via LC-MS/MS. Analogues that did not have calibration material were calibrated of the nearest structurally related analogue, as shown.

| **Toxin name** | **Abbreviation** | **Calibration standard** | **Calibrant** |
| --- | --- | --- | --- |
| Brevetoxin-1 | BTX-1 | N | BTX-2 |
| Brevetoxin-2 | BTX-2 | Y | BTX-2 |
| Brevetoxin-3 | BTX-3 | Y | BTX-3 |
| Brevetoxin-B1 | BTX-B1 | N | dBTX-B2 |
| Brevetoxin-B2 | BTX-B2 | N | dBTX-B2 |
| s-desoxybrevetoxin-B2 | dBTX-B2 | Y | dBTX-B2 |
| Brevetoxin-B4 | BTX-B4 | N | dBTX-B2 |
| Brevetoxin-B5 | BTX-B5 | Y | BTX-B5 |

Supplementary Information 1. Discussion of Morphological features and comparison with other *Karenia* reports

In examining the morphological features of the cultured strain and environmental samples (Fig. 3), they most closely resemble *Karenia cristata* Botes, Sym & Pitcher, because of the apical crest (for which it was named), its centrally located round nucleus, and the slightly asymmetrical hypocone (right lobe longer than left) (Fig. 3). While the type description^43^ specifies a centrally located nucleus, two other studies ^51 50^ describe the nucleus to be horizontally located in the hypocone. The SEM images of the South African type culture failed to resolve the sulcal intrusion, which was described as “closed” ^43^. In contrast our SEM images (Fig. 3) clearly show a wide-open open sulcal intrusion. The later described *Karenia hui* Lu ^16^ from Baguang Bay, China, has a similar apical crest but was differentiated from *K. cristata* by a wide-open sulcal intrusion and a horizontally elongated rather than circular nucleus in the centre of the hypocone. The species *K.hui* is highly morphologically similar to *K. cristata*, and differs in molecular genetic sequence by 22 bp over 883 bp of LSU rDNA (2.5 % seq. divergence). We established a new quantitative PCR assay specifically to detect *K. hui*, and found that it was not present in this HAB (Fig. 2).

Supplementary Fig. 1. Standard curves of species-specific assays for *Karenia* spp., presenting the quantification cycle (y-axis) versus the known cell number in log scale (x-axis). a, *Karenia brevisulcata* CAWD82, b, *Karenia longicanalis* CAWD457, c, *Karenia mikimotoi* CAWD05, d, *Karenia papilionacea* CAWD91, e, *Karenia cristata*, f, *Karenia hui* and g, *K. brevis* CAWD04

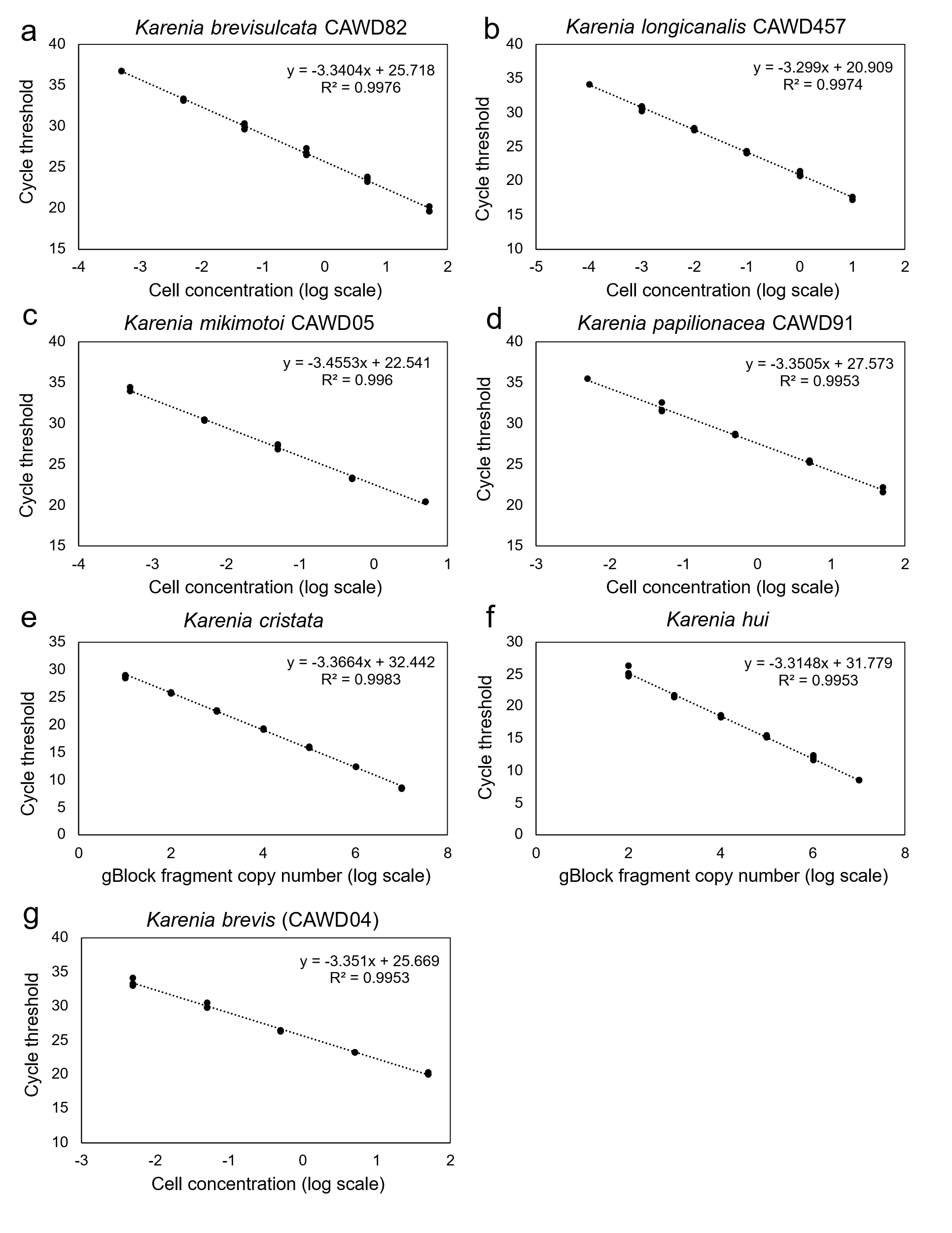

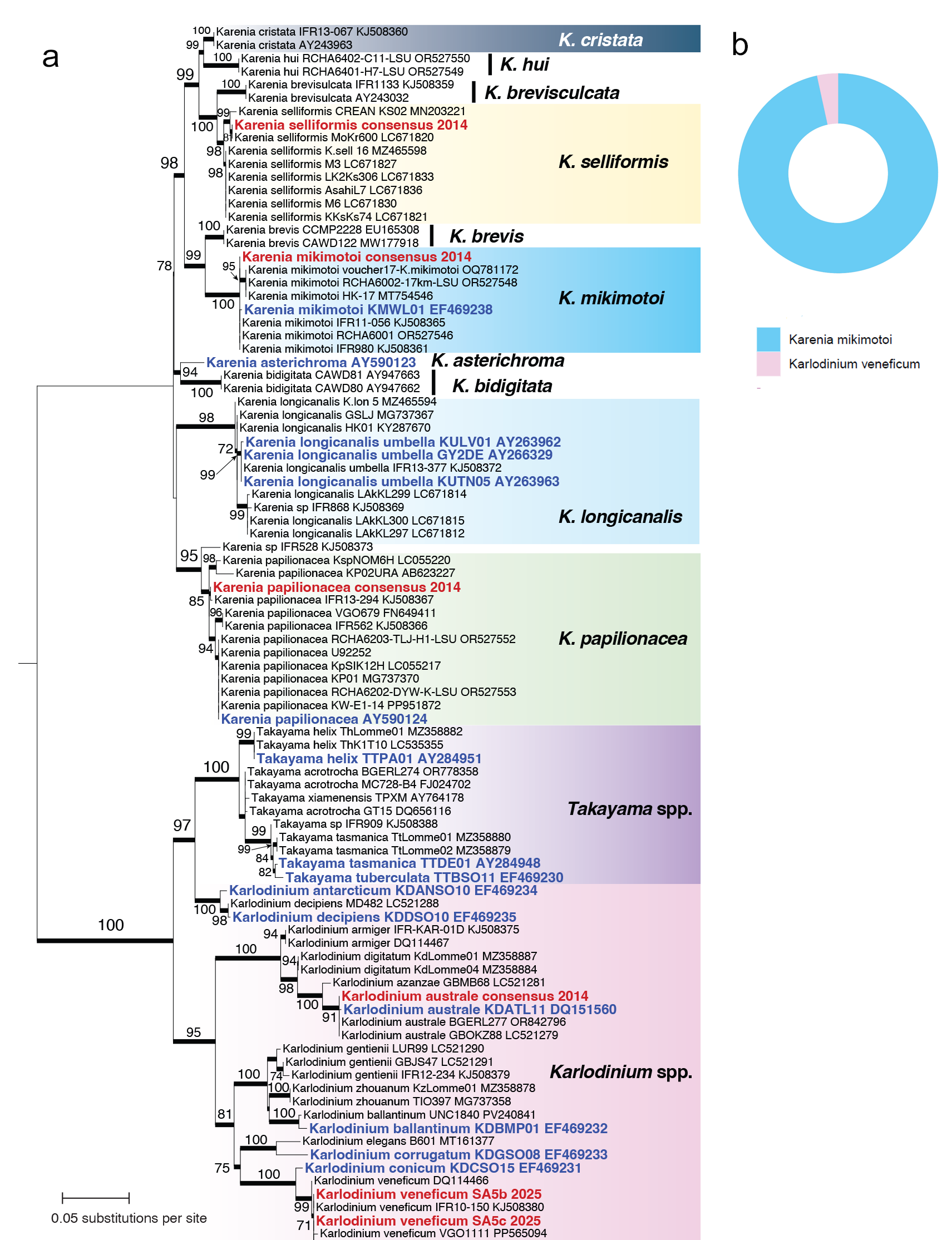

Supplementary Fig. 2. a. Maximum likelihood tree of Kareniaceae based on LSU rDNA sequences higlighting taxa detected in Australian waters prior to the 2025 bloom. The tree is rooted with the clade of *Takayama* and *Karlodinium* as outgroup. Re-analysed sequences from 2014 bloom in South Australia as well as sites in NSW in 2014 are highlighted in red. Sequences from taxa isolated in Australian waters are highlighted in blue. Ultrafast bootstrap support (>70%) is shown on each internal node, and a thick branch length indicates Bayesian posterior probability = 1.0 in the additionally inferred Bayesian tree (see Methods). Unit of branch length is number of substitutions per site .b. Donut plot of Kareniaceae ASVs from sample from the Coffin Bay (Fig 1a) *Karenia* HAB in 2014, showing it was mostly *K. mikimotoi* with *Karlodinium veneficum*.
